# The cryo-EM structure of ASK1 reveals an asymmetric architecture allosterically modulated by TRX1

**DOI:** 10.1101/2023.12.20.572539

**Authors:** Karolina Honzejkova, Dalibor Košek, Veronika Obsilova, Tomas Obsil

**Affiliations:** Department of Physical and Macromolecular Chemistry, Faculty of Science, Charles University, Prague, Czech Republic; Institute of Physiology of the Czech Academy of Sciences, Laboratory of Structural Biology of Signaling Proteins, Division BIOCEV, 252 50 Vestec, Czech Republic

**Author notes:** **Author Contributions:** T.O. and V.O. supervised the project and provided scientific guidance. K.H. performed protein expression/purification experiments, SV AUC measurements and data analysis and prepared samples for cryo-EM and HDX-MS measurements. D.K. performed cryo-EM measurements and data analysis, model building and structure refinement. K.H., D.K., V.O. and T.O. wrote the manuscript. All co-authors revised the manuscript.

**Keywords:** ASK1, MAP3K, MAPK signaling, thioredoxin

## Abstract

Apoptosis signal-regulating kinase 1 (ASK1) is a crucial stress sensor, directing cells towards apoptosis, differentiation and senescence via the p38 and JNK signaling pathways. ASK1 dysregulation has been associated with cancer and inflammatory, cardiovascular and neurodegenerative diseases, among others. However, our limited knowledge of the underlying structural mechanism of ASK1 regulation hampers our ability to target this member of the MAP3K protein family towards developing therapeutic interventions for these disorders. Nevertheless, as a multidomain Ser/Thr protein kinase, ASK1 is regulated by a complex mechanism involving dimerization and interactions with several other proteins, including thioredoxin 1 (TRX1). Thus, the present study aims at structurally characterizing ASK1 and its complex with TRX1 using several biophysical techniques. As shown by cryo-EM analysis, in a state close to its active form, ASK1 is a compact and asymmetric dimer, which enables extensive interdomain and interchain interactions. These interactions stabilize the active conformation of the ASK1 kinase domain. In turn, TRX1 functions as a negative allosteric effector of ASK1, modifying the structure of the TRX1-binding domain and changing its interaction with the tetratricopeptide repeats domain. Consequently, TRX1 reduces access to the activation segment of the kinase domain. Overall, our findings not only clarify the role of ASK1 dimerization and inter-domain contacts but also provide key mechanistic insights into its regulation, thereby highlighting the potential of ASK1 protein-protein interactions as targets for antiinflammatory therapy.

## Introduction

Mitogen-activated protein (MAP) kinase cascades are one of the most important signal transduction networks in cells. Highly conserved throughout eukaryotes^1^, MAP kinase pathways are activated in response to a number of stimuli, such as cytokines, growth factors, oxidative stress, calcium influx and lipopolysaccharides, promoting proliferation, inflammatory responses and apoptosis ^2^. The incoming signals are transmitted across a common, three-layered protein kinase system composed of the upstream MAP kinase kinase kinase (MAP3K), the intermediate MAP kinase kinase (MAP2K) and the downstream MAP kinase (MAPK)^1^. MAP2Ks and MAPKs are similarly activated through phosphorylation by an upstream kinase, but MAP3K activation is much more complex. MAP3Ks require a tight regulation given the high number of their potential triggers and the serious consequences of their dysregulation, in the form of various diseases^3,4^.

Cancer and inflammatory, cardiovascular, and neurodegenerative diseases, among others, have been extensively associated with excessive apoptosis signal-regulating kinase 1 (ASK1) signaling, in particular^5-8^. Also known as MAP3K5, ASK1 stands out for its role in directing cells towards apoptosis, differentiation and senescence via the p38 and JNK MAP kinase pathways^9,10^. However, currently available JNK and p38 inhibitors lack efficacy and/ or have undesirable side effects^11,12^. Therefore, as their upstream activator, ASK1 is a prospective target for therapeutic intervention in these disorders, especially considering its wide range of triggers and interactions.

Various stress stimuli, such as oxidative and endoplasmic reticulum (ER) stress and calcium influx, activate ASK1, thus explaining why this Ser/Thr-specific protein kinase is such a crucial stress sensor^9^. In line with this function, human ASK1 is a multi-domain protein consisting of an N-terminal thioredoxin-binding domain (TBD), a central regulatory region (CRR) formed by tetratricopeptide repeats (TPR) and pleckstrin-homology (PH) domains, a kinase domain (KD) and a C-terminal coiled-coil (CC) region followed by a sterile alpha motif (SAM) domain (Fig. 1a)^13-16^. In both resting and active states, ASK1 forms a large multiprotein complex, the ASK signalosome, with dozens of interacting proteins, including other ASK family members ASK2 and ASK3^17-20^. In the resting state, ASK1 constitutively oligomerizes, presumably via the C-terminal CC region and the SAM domain, remaining inactive through interactions with thioredoxin 1 (TRX1), glutaredoxin and 14-3-3 proteins^3,13,15-18,21-24^.

**Figure 1.**
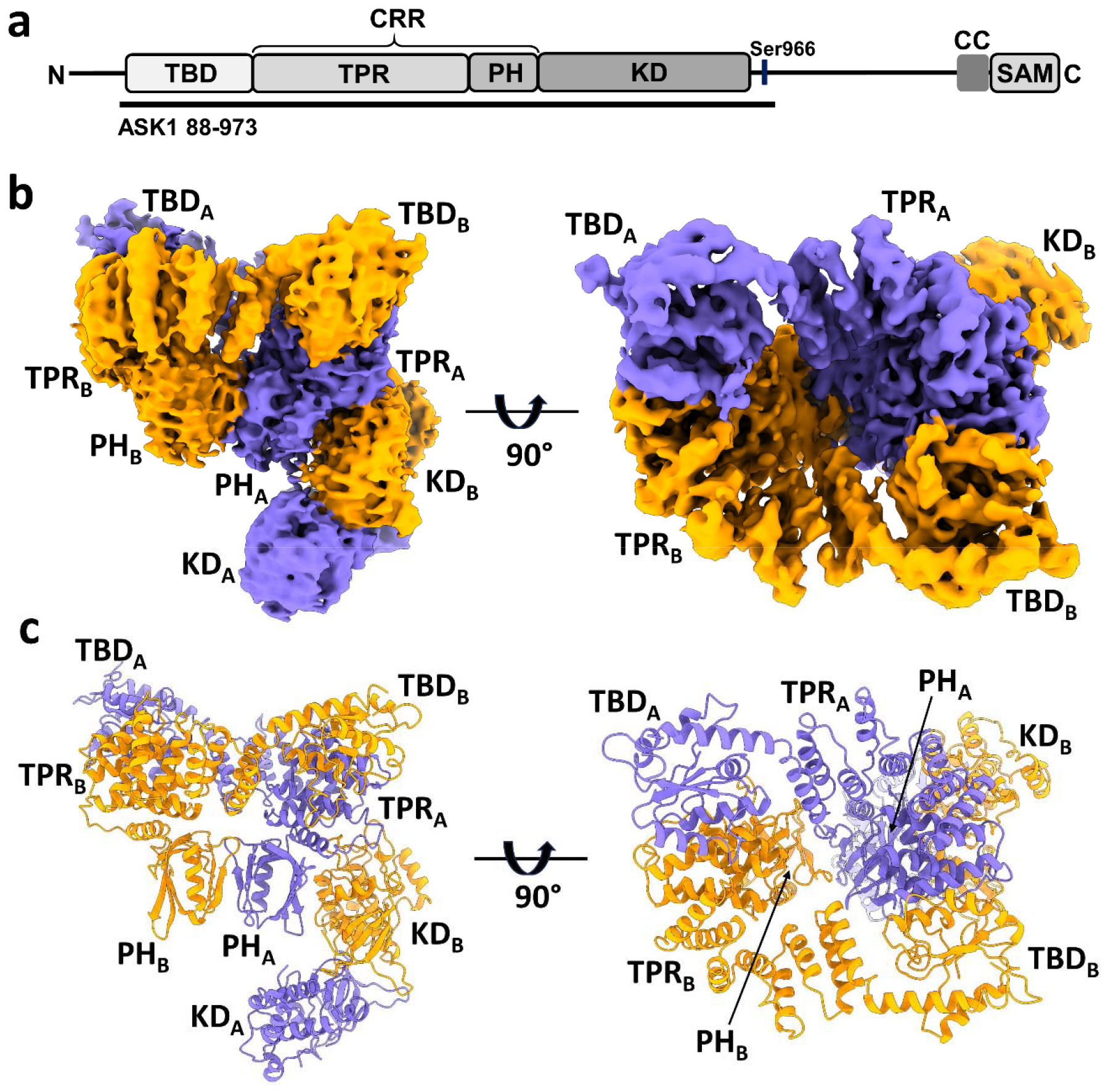
Structure of C-terminally truncated ASK1. (a) Schematic domain structure of ASK1. TBD, thioredoxin-binding domain; TPR, tetratricopeptide repeats; PH, pleckstrin-homology domain; CRR, central regulatory region; KD, kinase domain; CC, coiled-coil motif; SAM, sterile alpha motif domain. Black bar represents construct used in cryo-EM analysis. (b, c) Cryo-EM density map and cartoon view of C-terminally truncated dimeric ASK1. Density maps were generated using threshold level 5.5.

Although the exact role of these binding partners in ASK1 regulation is still debated, TRX1 binding to the N-terminal TBD is thought to prevent homophilic interactions between ASK1 N-termini containing the TBD, TPR, PH and KD domains^3,14^. When complexed with TRX1, inactive ASK1 is ubiquitinated and degraded^25^. By contrast, under oxidative stress, TRX1 oxidation triggers its dissociation from the ASK signalosome, followed by 14-3-3 dissociation and tumor necrosis factor receptor-associated factor (TRAF2/5/6) recruitment to CRR^3,15,26-33^. Presumably through CRR or another ASK1 domain N-terminal to KD, death domain-associated protein 6 (Daxx) directly interacts with ASK1, thereby mediating ASK1 activation by the death-inducing ligand system, known as cluster of differentiation 95 (CD95)/Fas receptor^34^. These events likely enable homophilic interactions between ASK1 N-termini, leading to Thr838 phosphorylation in the activation segment either by trans-autophosphorylation or by protein serine/threonine kinase 38 (MPK38) and subsequent ASK1 activation^21,22,31,35^.

Combined, these findings have advanced our knowledge of the structure of individual ASK1 domains^13-16^. Yet, several questions as to how these domains interact with each other, how they participate in ASK1 oligomerization, or what role these interactions play in ASK1 regulation remain unanswered. Moreover, the mechanism whereby TRX1 binding to the N-terminal TBD inhibits ASK1 activation is also unresolved. CRR may keep TBD and KD relatively close, enabling TRX1 to block CRR and/or KD and, hence, preventing substrate binding and inhibiting ASK1^14^. But ASK1 dimerization most likely significantly affects not only the conformation of ASK1 but also its interdomain interactions. Considering the above, overcoming current obstacles to the development of effective drug therapies targeting ASK1 requires acquiring relevant structural data to further our understanding of the complex regulation of ASK1.

To gain structural insights into ASK1 regulation and the role of TRX1 binding, this study aims at structurally characterizing the dimeric, C-terminally truncated ASK1 with the domains TBD, TPR, PH and KD in a state close to its active form and its complex with TRX1. For this purpose, we used several biophysical methods, including cryo-electron microscopy (cryo-EM), hydrogen/deuterium exchange coupled to mass spectrometry (HDX-MS) and sedimentation velocity analytical ultracentrifugation (SV AUC). Our data reveal that ASK1 forms a compact and asymmetric dimer in which all four N-terminal domains are involved in extensive interdomain and interchain interactions. These interactions stabilize the active conformation of ASK1 kinase domain. TRX1 functions as a negative allosteric effector of ASK1 by modulating the structure of all its N-terminal domains, including the activation segment of the catalytic domain. Overall, our findings not only clarify the role of ASK1 dimerization and inter-domain contacts but also provide key mechanistic insights into its regulation.

## Results

### ASK1 forms an asymmetric and compact dimer through extensive inter-domain and inter-chain interactions

Given the low expression yield and solubility of full-length human ASK1, we designed a C-terminally truncated construct consisting of TBD, CRR and KD (residues 88-973), thus all domains crucial for ASK1 regulation. The expression yield and stability of this protein was sufficient for subsequent studies. Cryo-EM imaging of the ASK1 TBD-CRR-KD revealed well-dispersed particles, with 2D class averages showing obvious secondary structure elements (Supplementary Fig. S1). Approximately 5,780 micrograph movies enabled single-particle reconstructions of this protein at a nominal resolution of 3.7 Å, as further detailed in Methods and Supplementary Table S1 and Fig. S2.

The cryo-EM map revealed that C-terminally truncated ASK1 forms a compact and asymmetric dimer, enabling extensive interdomain and interchain interactions (Fig. 1b,c and Supplementary Fig. S3). One side of the molecule is formed by TBD and TPR domains of both protomers, with TBD domains embedded between TPR domains. The TPR domains then interact with a dimer of PH domains, and the resulting dimeric TBD-TPR-PH module has a funnel shape with approximately 2-fold rotational symmetry. The other side of the molecule is formed by a KD dimer, which interacts with the N-terminal TBD-TPR-PH module through the KD of only one protomer. The binding site of this KD is located at the interface between the TPR and PH domains of the opposite protomer, and the second KD has no contacts with the N-terminal domains of either its own or the other chain.

### All N-terminal domains participate in ASK1 homodimerization

The region of the cryo-EM map that corresponds to TBD domains (residues 95-266) was interpreted using the AlphaFold model of ASK1 (AF-Q99683-F1). This model suggested that TBD consists of a six-stranded, mostly parallel, β-sheet decorated with several α-helices, thus resembling the thioredoxin structure. Such a conformation was in line with our cryo-EM map, which revealed a similar arrangement of secondary structure elements in both TBDs (Fig. 2a and Supplementary Fig. S4). TBD is a compact and globular domain that interacts with the TPR domain of the opposite protomer via the helix α3 and the loop between α4 and β5. The C-terminal helix α5 of TBD is connected by a short loop to helix α6, which is the first helix of the TPR domain. In the same chain, there are no other contacts between TBD and TPR except for the interaction between the C-terminus of helix α3 of TBD and the N-terminus of helix α6 of TPR.

**Figure 2.**
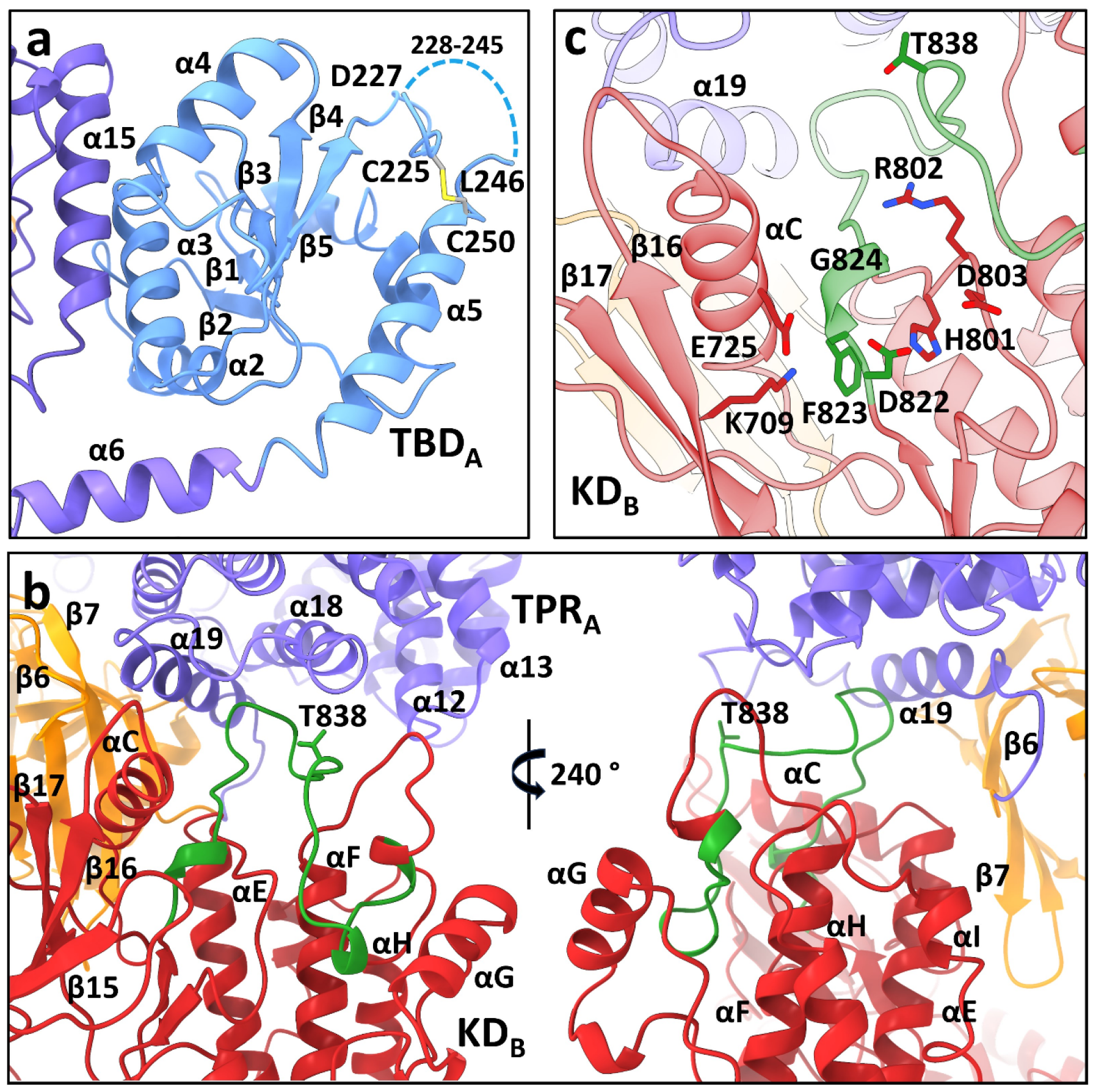
All four N-terminal domains of ASK1 are involved in extensive interdomain and interchain interactions. (a) Cartoon representation of TBD_A_ and its interaction with helix α15 of TPR_B_. The dashed line indicates the missing section (residues 228-245). (b) Cartoon representation of interactions between KD_B_, TPR_A_ and PH_A_ domains (shown in red, violet and orange, respectively). The activation segment is shown in green. (c) Detailed view of the active site of KD_B_. KD_B_ is shown in red, TPR_A_ is shown in violet. The activation segment is shown in green.

In a previous study, cysteine C250, located at the N-terminus of α5, was identified as a crucial residue for TBD structural integrity and TRX1 binding^15,23^. Corroborating these findings, our structure of TBD_A_ suggests that this cysteine residue forms a disulfide bridge with cysteine C225 located nearby (Fig. 2a). This disulfide, whose formation was shown previously^33^, likely stabilizes the region containing residues 216-245 located between β5 and α5. The quality of the cryo-EM map in this region is worse for TBD of the opposite chain (TBD_B_), but here too C225 and C250 are close to each other. In addition, this region appears to be very flexible as no interpretable density was found for residues 228-245 in both TBDs. This result is consistent with a previous NMR characterization of isolated TBD, which showed that this domain retains substantial conformational plasticity^15^.

The conformation of both CRR domains resembles the crystal structure of isolated CRR, with 14 tightly arranged α-helices (α6-α19) forming seven tetratricopeptide repeats (TPRs), followed by a PH domain^14^ (Fig. 1c and Supplementary Fig. S4). Superpositions with the crystal structure showed minor helix shifts in the first four repeats of both TPRs, most likely due to interactions with the TBD of the opposite chain (Supplementary Fig. S5). Both CRRs interact directly only through the β-sheets of their PH domains, and the resulting CRR dimer has 2-fold rotational symmetry (Fig. 1b,c). Combined, extensive interactions between N-terminal TBD, TPR and PH domains emerge as a crucial factor for the homodimerization of C-terminally truncated ASK1.

### Interdomain interactions stabilize the activation segment of the kinase domain

TPR and PH domains of one chain (chain A) create a docking platform for the KD of the opposite chain (KD_B_), which interacts with TPRs 4 (helices α12, α13) and 7 (helices α18, α19) of TPR_A_, and β6 and β7 strands of PH_A_, mainly via its C-lobe (Fig. 2b). The kinase domains of both protomers dimerize, as previously observed in the crystal structure of isolated ASK1 KD with bound inhibitor staurosporine ^13^, i.e., in a head-to-tail orientation through an extensive interface spanning almost the entire length of the domain. However, its comparison with the crystal structure showed changes in the position of the αC (α21) helix, which is in the inward active position, in both KDs (Supplementary Fig. S6a,b). This shift positions the conserved active site lysine (K709) within hydrogen-bond distance of the αC glutamate (E725), as usually found in active kinases^36^ (Fig. 2c and Supplementary Fig. S6e).

The cryo-EM map of KD_B_, which interacts with TPR_A_ and PH_A_ domains, enabled us to build the whole activation segment (residues 822-849). This segment adopts a conformation competent for substrate binding (Fig. 2b). A similar conformation was observed in the crystal structure of ASK1 KD, which, however, showed only the beginning and end of the activation segment (Supplementary Fig. S6b)^13^. In this conformation, the conserved residue E725 from the αC (α21) helix is within hydrogen-bond distance of the main-chain of the DFG motif residues F823 and G824 at the beginning of the activation segment (Fig. 2c and Supplementary Fig. S6e).

A similar conformation of the activation segment was also observed in the active form of another MAP3K BRAF (Supplementary Fig. S6c)^37^. In active BRAF, the conformation of the activation segment is stabilized by E611 interactions with R^575^ from the catalytic HRD motif, but in the KD_B_ of the present structure, the activation segment is stabilized by interactions with the α18 and α19 helices from the last TPR repeat of the TPR_A_ domain (Fig. 2b).

The activation segment of the KD domain from the opposite chain (KD_A_), which has no contacts with N-terminal domains, is not visible in our map, likely due to its high flexibility. However, the beginning and end of this segment adopt a conformation similar to that observed in the crystal structure of the isolated ASK1 KD^13^. Together with the inward position of the helix αC (α21), this conformation suggests that KD_A_ is also in an active conformation (Supplementary Fig. S6a).

The comparison with the crystal structure of isolated KD revealed a change in the relative position of the KD subunits. The superimposition of the KD dimer of ASK1 TBD-CRR-KD with the crystal structure of the KD dimer ^13^ through KD_A_ is shown in Supplementary Fig. S6d. KD_B_ is slightly rotated relative to the equatorial plane of the dimer due to interactions of its C-lobe with TPR_A_. This rotation results in a ∼4 Å shift of the C-lobe relative to its position in the crystal structure of the KD dimer. Overall, interdomain interactions within ASK1 TBD-CRR-KD stabilize the conformation of the activation segment. As a result, the activation segment remains competent for substrate binding.

### TBD-CRR, but neither TBD nor CRR form dimers in solution

Previous studies have shown that isolated KD forms a stable dimer with a *K*_D_ of ∼220 nM in which the protomers are oriented in a head-to-tail manner and interact through a large dimerization interface^13,24^. To verify the interdomain contacts observed in the cryo-EM map and their relevance to the homodimerization of C-terminally truncated ASK1, we prepared isolated TBD, CRR and KD and two additional constructs composed of TBD-CRR and CRR-KD to study their oligomerization by sedimentation velocity analytical ultracentrifugation (SV AUC).

The sedimentation coefficient distribution *c*(*s*) of the isolated ASK1 KD assessed by SV AUC showed only one peak with weight-average sedimentation coefficients corrected to 20.0 °C and to the density of water, *s*_w(20,w)_, of 4.2 S (estimated *M*_w_ ∼62 kDa), most likely corresponding to the KD dimer (the theoretical *M*_w_ of the KD dimer is 72.4 kDa) (Fig. 3c). In contrast, the isolated domains, TBD and CRR, were protomeric in solution as their sedimentation coefficient distributions *c*(*s*) showed peaks with *s*_w(20,w)_ of 2.2 and 3.3 S with estimated *M*_w_ ∼16 kDa and ∼38 kDa, respectively (theoretical *M*_w_ of TBD and CRR are 20.6 and 45.5 kDa, respectively) (Fig. 3a,b). While the ASK1 TBD-CRR construct showed concentration-dependent dimerization, as indicated by its bimodal *c*(*s*) distribution (Fig. 3d), the ASK1 CRR-KD construct formed a stable dimer in solution, with a *s*_w(20,w)_ of 6.9 S (estimated *M*_w_ ∼166 kDa, theoretical *M*_w_ 163 kDa) (Fig. 3d,e). Similarly, as shown by its *c*(*s*) distribution, the longest ASK1 TBD-CRR-KD construct also formed stable dimers in solution, with a *s*_w(20,w)_ of 8.5 S (Fig. 3f). This value is consistent with the theoretical sedimentation coefficient value of 9 S calculated using the HydroPRO program^38^ and the cryo-EM structure of the ASK1 TBD-CRR-KD dimer. The *c*(*s*) distribution of ASK1 TBD-CRR-KD also revealed higher oligomers, whose abundance increased with the concentration.

**Figure 3.**
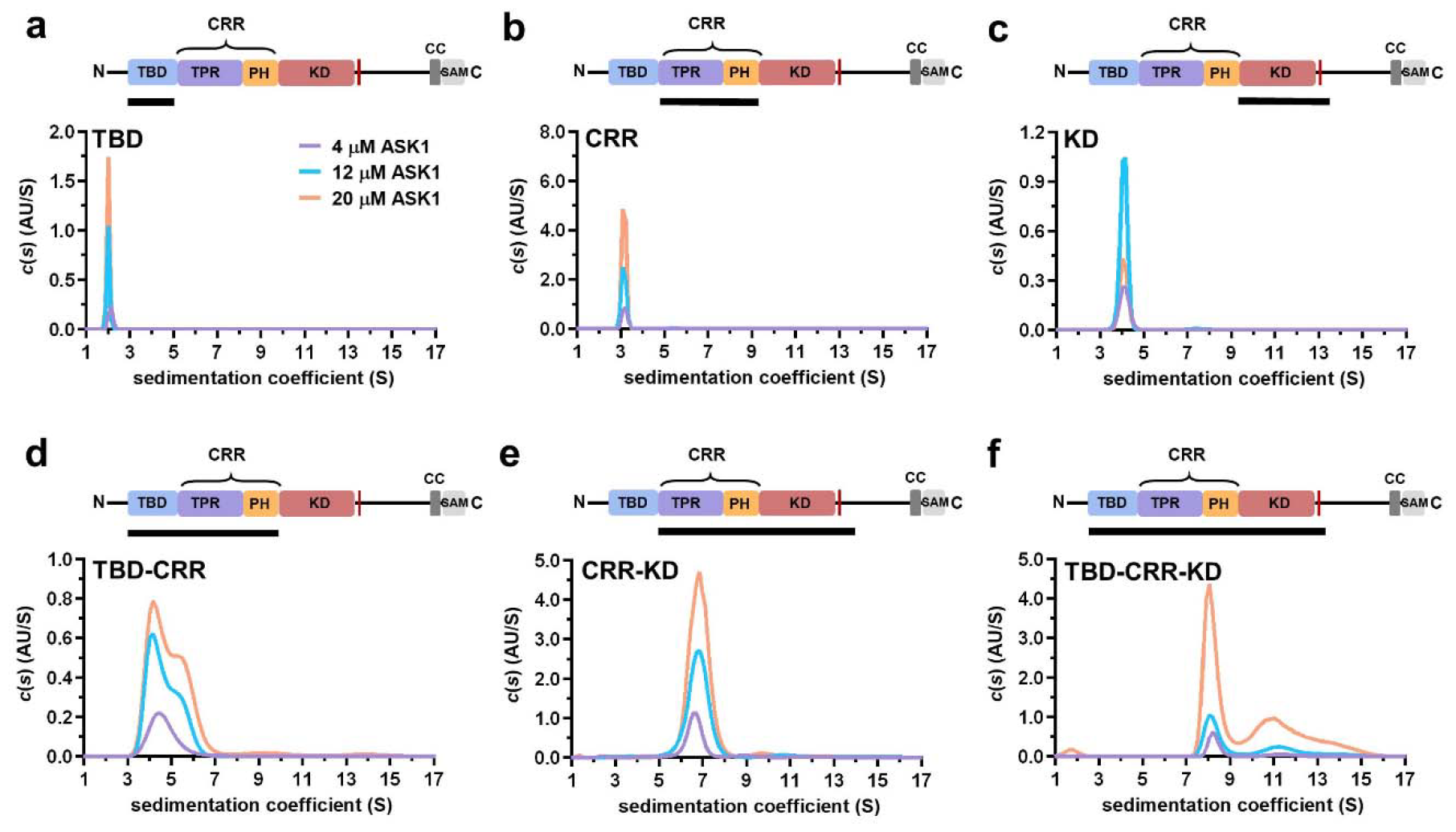
TBD-CRR forms dimers in solution, but not TBD or CRR. (a-f) Sedimentation coefficient distributions (*c*(*s*)) of different ASK1 N-terminal constructs (TBD, CRR, KD, TBD-CRR, CRR-KD and TBD-CRR-KD) measured at concentrations of 4 μM, 12 μM and 20 μM. Schematic domain structure of ASK1 is shown at the top; the black bar represents the used construct. TBD, thioredoxin-binding domain; TPR, tetratricopeptide repeats; PH, pleckstrin-homology domain; CRR, central regulatory region; KD, kinase domain; SAM, sterile alpha motif domain.

The dimerization of KD-containing ASK1 constructs is not surprising because KDs form stable dimers, and CRRs do not interfere with the dimerization interface; on the contrary, they facilitate dimerization through interactions between PH domains (Fig. 1b,c). Furthermore, the absence of dimerization of an isolated TBD is consistent with its interactions within the N-terminal TBD-CRR module, wherein TBDs interact only with TPR, not with each other. Isolated CRR showed no dimerization either, suggesting that contacts between PH domains observed in the ASK1 TBD-CRR-KD dimer are not strong enough to allow the formation of a stable CRR dimer and likely result from KD dimerization. Combined, our SV AUC measurements with isolated domains and their pairs are consistent with the cryo-EM structure of C-terminally truncated ASK1 and confirm that all three N-terminal domains are involved in homophilic interactions between ASK1 chains.

### TRX1 binding induces structural changes in the activation segment of ASK1 KD

TRX1 binding presumably inhibits ASK1 activation by blocking the homophilic interaction of the N-terminal part of ASK1, according to the currently accepted model of ASK1 activation under oxidative stress conditions^3,18^. Since we were unable to prepare a sufficiently stable ASK1:TRX1 complex for cryo-EM analysis, apparently due to their relatively weak interactions under the conditions used in this study^23^, we assessed TRX1 binding effects on ASK1 TBD-CRR-KD structure and dimerization by SV AUC and HDX-MS. Unexpectedly, SV AUC showed that TRX1 binding has no effect on ASK1 TBD-CRR-KD dimerization. However, when we compared *c*(*s*) distributions, we found a significant reduction in peak area, in the region of sedimentation coefficients 10-12 S, in the presence of TRX1. This result indicates that TRX1 prevents the formation of higher oligomers (Supplementary Fig. S7).

By HDX-MS, we monitored the kinetics of hydrogen-to-deuterium exchange along the polypeptide backbone because this method enables us to evaluate the structure of proteins and their complexes^39^. Hydrogens of amide groups involved in stable hydrogen bonds and/or sterically shielded from the solvent are protected from exchange. In contrast, flexible regions exposed to the solvent exchange more quickly than rigid and buried regions.

The comparison of ASK1 TBD-CRR-KD deuteration profiles with and without TRX1 revealed that several TBD regions were significantly (more than twice the standard deviation and above 3%) less deuterated, that is, more protected, upon TRX1 binding (Fig. 4a,b and Supplementary Figs. S8-S10). More specifically, significant protection was observed in residues 128-143, 184-195, and especially in segment 206-259. Many of these segments overlapped with regions previously identified by NMR as the TRX1 binding surface of TBD^15^ and, moreover, included the flexible region between β5 and α5 containing C225 and C250 (Fig. 4b and Supplementary Fig. S9). These regions on the flexible outer surface of the TBD near the interface between the TBD and the TPR of the opposite chain may form a TRX1-binding site.

**Figure 4.**
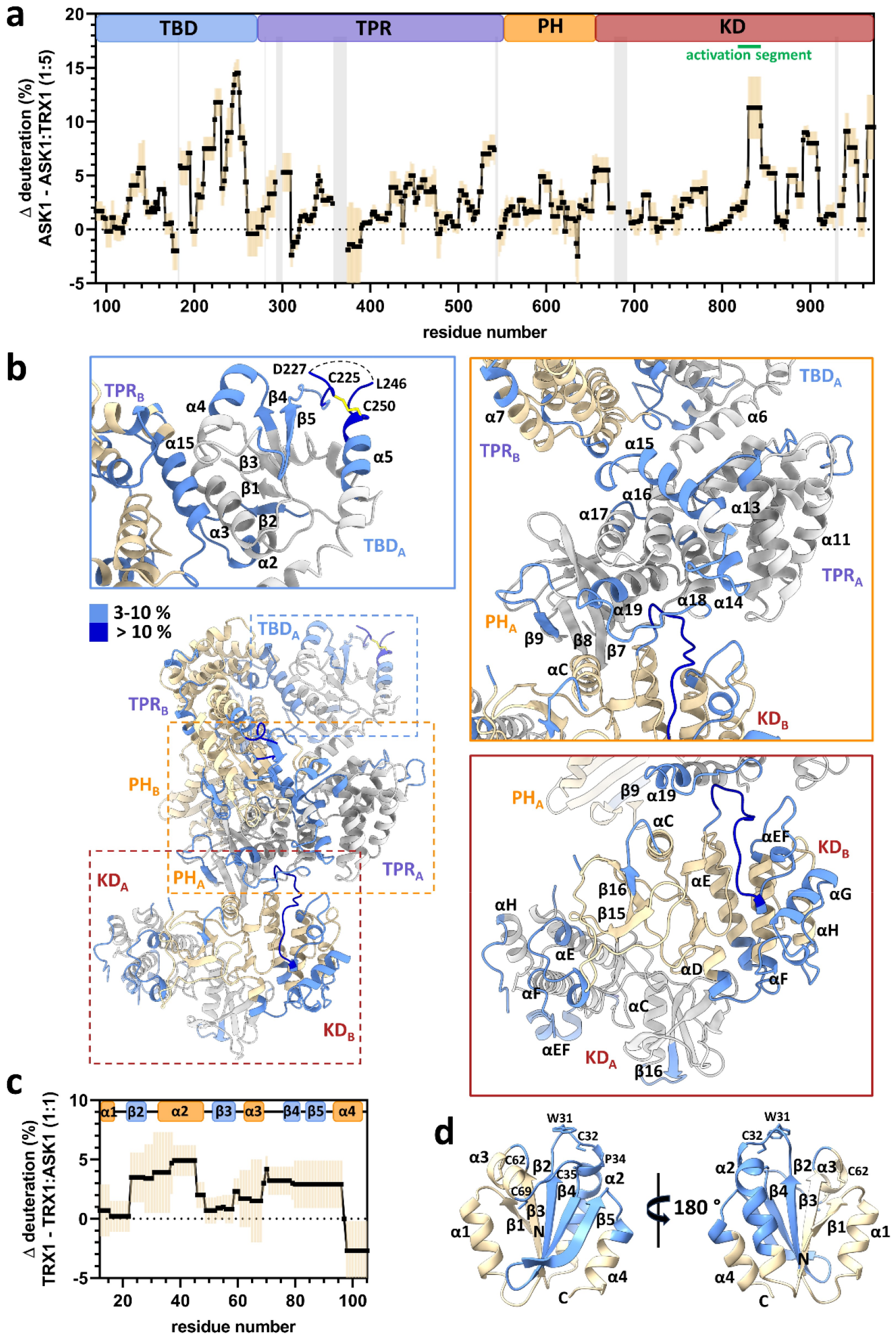
TRX1 binding induces structural changes in all N-terminal domains of ASK1. (a) Differences in ASK1 TBD-CRR-KD deuteration with and without TRX1 after 600 s. Positive values indicate protection (lower deuterium uptake) after TRX1 binding. The graph shows the average of three replicates (black points) and the standard deviation (light orange). Grey zones indicate areas without coverage. The domain structure of ASK1 TBD-CRR-KD is shown at the top. TBD, thioredoxin-binding domain; TPR, tetratricopeptide repeats; PH, pleckstrin-homology domain; CRR, central regulatory region; KD, kinase domain. (b) Cartoon representation of the structure of the ASK1 TBD-CRR-KD dimer (chain A in white, chain B in light yellow) colored according to changes in deuteration in the presence of TRX1 after 600 s. Changes in deuteration greater than twice the standard deviation and above 3% were considered significant. The insets show detailed views of the TBD_A_ and its interface with TPR_B_ (blue box), CRR_A_ (orange box) and KD dimer (brown box). (c) Differences in TRX1 deuteration with and without ASK1 TBD-CRR-KD after 600 s. Positive values indicate protection (lower deuterium uptake) after ASK1 TBD-CRR-KD binding. The graph shows the average of three replicates (black points) and the standard deviation (light orange). The secondary structure of TRX1 is shown at the top. (d) Cartoon representation of TRX1 structure (in reduced state, PDB ID: 1ERT^52^) colored according to changes in deuteration after ASK1 TBD-CRR-KD binding for 600 s.

This hypothesis is in line with the decrease in deuteration observed in the α15 helix of the TPR domain, which interfaces with TBD, in the adjacent α13, α14 and α19 helixes, and in the α13-α14, α14-α15, α17-α18 and α18-α19 loops (Fig. 4a,b and Supplementary Fig. S11). Within the PH domain, the β-strand β9 and the β8-β9 loop, located near the α19 helix of the TPR, were also protected. Through interactions, these regions connect the TRX1-binding site of TBD with the TPR and PH regions that form the docking surface of one of the two KDs (Supplementary Fig. S9). Given the lower deuteration of these regions, TRX1 binding likely stabilizes their structure and/or reduces their solvent accessibility (e.g., through conformational change), as shown by changes in the deuteration of several KD regions.

TRX1 binding to TBD reduced deuteration near the active site and at the C-terminus of the KD (Fig. 4a,b and Supplementary Fig. S12). The most protected area included residues 830-844, which form the activation segment and interact with the TPR domain. In addition to the activation segment, the region containing αEF (α24), the helix αG (α26) and the αD-αE (α22-α23), αF-αG (α25-α26) and αG-αH (α26-α26) loops also showed reduced deuteration. Accordingly, these regions likely interact with each other, forming a coherent area whose structure and/or access to solvent is affected by TRX1 binding.

Significant protection was also observed in residues 655-670, which form a linker connecting PH and KD (Fig. 4a). This linker should be quite flexible as no interpretable density was found in this region, in either protomer. However, TRX1 binding reduced its deuteration, suggesting structural changes at the interface of PH and KD. Taken together, our HDX-MS results indicate that TRX1 binding to TBD significantly alters the interactions and structure of TBDs, leading to changes in interactions within CRRs and to conformational changes in KDs, including in their activation segment and the C-terminus.

### TRX1 interacts with ASK1 through its active site

Based on comparisons of TRX1 deuteration profiles with and without ASK1 TBD-CRR-KD, the helix α2 region and the β-strands β1, β3, and β4 (Figs. 4c,d and Supplementary Fig. S13 and S14) are significantly protected. This protection includes the highly conserved catalytic motif W31CGPC35, located at the N-terminus of the α2 helix, whose sulfhydryl groups are responsible for TRX-dependent redox activity^40^. This result is ccording to which TRX1 binds to ASK1 through its active _site_3,15,18,33.

## Discussion

Homophilic interactions between N-terminal domains are crucial for ASK1 activation, as shown by our structural analysis of the C-terminally truncated ASK1. In addition to these interactions, ASK1 oligomerization is mediated by its C-terminal CC motif because the C-terminally truncated ASK1 has lower Thr838 phosphorylation levels and basal activity^22^. Furthermore, the ASK1 construct with the first 384 residues of this protein, containing TBD and the N-terminal portion of TPR, can also oligomerize, as previously demonstrated by co-immunoprecipitation ^3^ and in line with our SV AUC measurements (Fig. 3d) and with our cryo-EM analysis of the apo form of ASK1 TBD-CRR-KD. This analysis revealed a compact and asymmetric dimer with all four N-terminal domains involved in extensive interdomain and interchain interactions. These interactions stabilize the active conformation of the kinase domain of ASK1 (Figs. 1 and 2), highlighting their importance for ASK1 activation.

When comparing the conformation of KDs of the ASK1 TBD-CRR-KD dimer with the crystal structure of the isolated KD with bound inhibitor^13^, we noted a shift in the αC helix, occupying an inward position typical of the active conformation (Supplementary Fig. S6a,b). Considering this position and the position of the activation segment, which is structured through interactions with the adjacent TPR and maintains a conformation competent for substrate binding, in the case of KD_B_ (Fig. 2b,c), interactions with the dimeric N-terminal TBD-CRR module may stabilize the active conformation of one of the two KDs. Furthermore, located in the activation segment and known to play a key role in ASK1 activation^22^, T838 is oriented toward the solvent, so its (auto)phosphorylation could further stabilize the activation segment through interactions with residues (e.g. K526) of the loop between α18 and α19 of TPR located nearby.

In a recent study, we have shown that ASK1 TBD is structurally heterogeneous and has a globular conformation resembling the thioredoxin fold, with its C-terminal half (residues ∼165-260) forming a TRX1-binding site^15^. In this study, our cryo-EM structure revealed that these residues form not only the solvent-accessible outer surface of the ASK1 dimer but also the binding interface with the TPR domain (Fig. 2a). Consequently, TRX1 binding significantly affects the structure of TBD and its interactions within the ASK1 dimer.

Our HDX-MS measurements after TRX1 binding revealed considerable changes in deuteration kinetics, in three regions of TBD, including its C-terminal half. TRX1 binding also affected the deuteration kinetics of several regions of the TPR and PH domains of CRR (Fig. 4a,b and Supplementary Figs. S9-S11), suggesting conformational changes, presumably a shift or change in the relative orientation of the TPR repeats, in this part of ASK1^41^. Considering the differences between the TPR of the present ASK1 structure and the crystal structure of CRR (Supplementary Fig. S5)^14^ and its position in the ASK1 dimer, the ASK1 TPR domain appears to be a flexible structure. This structural flexibility enables communication between TBD and KD, as evidenced by structural changes in KD induced by TRX1 binding and is in line with the role of TPR domains as structurally diverse scaffolding and binding modules, some of which are quite flexible, triggering allosteric effects and supporting molecular switch functions^42,43^. For example, allosteric effects regulate the binding of Hif, a TPR protein, to histone complexes H2A-H2B and H3-H4^44^. Moreover, the flexibility or conformational plasticity of CRR may be involved in substrate recruitment^14^.

Many protein kinases are regulated through phosphorylation of a residue(s) located in the activation segment^36^. For example, Akt1 kinase is autoinhibited through intramolecular interactions between its PH and KD domains, limiting access to the activation segment^45^. Our HDX-MS data suggest that ASK1 is regulated by a similar mechanism involving structural changes in the activation segment. As evidenced by the slower deuteration kinetics, TRX1 substantially reduces solvent access to this segment and/or decreases its flexibility by altering interactions either within KD and/or with TPR (Fig. 4a,b), which may influence its phosphorylation on T838 within the signalosome. In doing so, TRX1 likely contributes to ASK1 inhibition^46,47^. Therefore, limiting access to residues of the activation segment regulated by phosphorylation may be thus a key regulatory event of ASK1 function.

In addition to the activation segment, TRX1 binding also decreased deuteration kinetics in other regions of KD, including the αG helix and surrounding loops (Fig. 4a,b). These regions are adjacent to the activation segment, and in MAP3K BRAF, the helix αG is involved in binding to its substrate MEK1^37^. Accordingly, TRX1 may function as an allosteric effector, changing the conformation of key regions of the KD domain when binding to TBDs and, therefore, inducing conformational changes transferred via CRRs.

Rather than disrupting homophilic interactions of the N-terminal domain, TRX1 binding alters them, since no significant deprotection was detected by HDX-MS. These results were also consistent with SV AUC measurements. However, at intracellular ASK1 concentrations, which are substantially lower than those used in our SV AUC and HDX-MS experiments, TRX1 binding may reduce homophilic interactions of the N-terminal domains of ASK1, as previously shown by immunoprecipitation analysis^3^. The resulting shift in the equilibrium toward the protomer may explain this difference between our SV AUC and HDX-MS measurements and previously reported findings.

Inactive ASK1 is bound to scaffolding 14-3-3 proteins, which recognize a motif containing phosphorylated S966, located at the C-terminus of KD^29,48^. 14-3-3 proteins suppress the catalytic activity of ASK1 through an unknown mechanism, albeit potentially involving suppression of homophilic interactions between N-terminal domains. In contrast, ASK1 activation requires TRAF2/5/6 or Daxx recruitment. These proteins interact with CRR and enhance N-terminal homophilic interactions of ASK1 protomers^3,26,27,34^. Thus, TRAF2/5/6 or Daxx binding to CRR could stabilize the conformation of the ASK1 dimer, thereby promoting its activation.

Our structural analysis was performed with a C-terminally truncated ASK1 missing the CC motif and a C-terminal SAM domain, involved in ASK1 oligomerization. Thus, we cannot rule out the possibility that C-terminal segments may affect interactions of N-terminal domains, including KDs. In addition, the structures of relevant complexes must be solved in subsequent studies to elucidate in detail structural changes caused by TRX1 binding, as well as other binding partners, and to fully understand how these interactions contribute to ASK1 regulation. Notwithstanding these limitations, our data provide the first structural insights into C-terminally truncated ASK1 in a state close to its active form. ASK1 forms a compact and asymmetric dimer with all four N-terminal domains involved in extensive interdomain and interchain interactions that stabilize the active conformation of ASK1 KD. TRX1, a negative regulator of ASK1, functions as an allosteric effector. TRX1 binding affects the structure of TBD and its interaction with TPR, thereby affecting the structure of CRR and allosterically modulating KD, even reducing access to the activation segment with the key phosphorylation site T838 (Fig. 5). Therefore, our findings open up opportunities for targeting (the) interaction(s) responsible for ASK1 activation towards developing selective ASK1 signaling inhibitors and ultimately pharmaceutical drugs for several inflammatory, cardiovascular and neurodegenerative diseases, among others^49,50^. Moreover, these results should prompt further research on this key MAP3K and its regulation, with a significant translational output.

**Figure 5.**
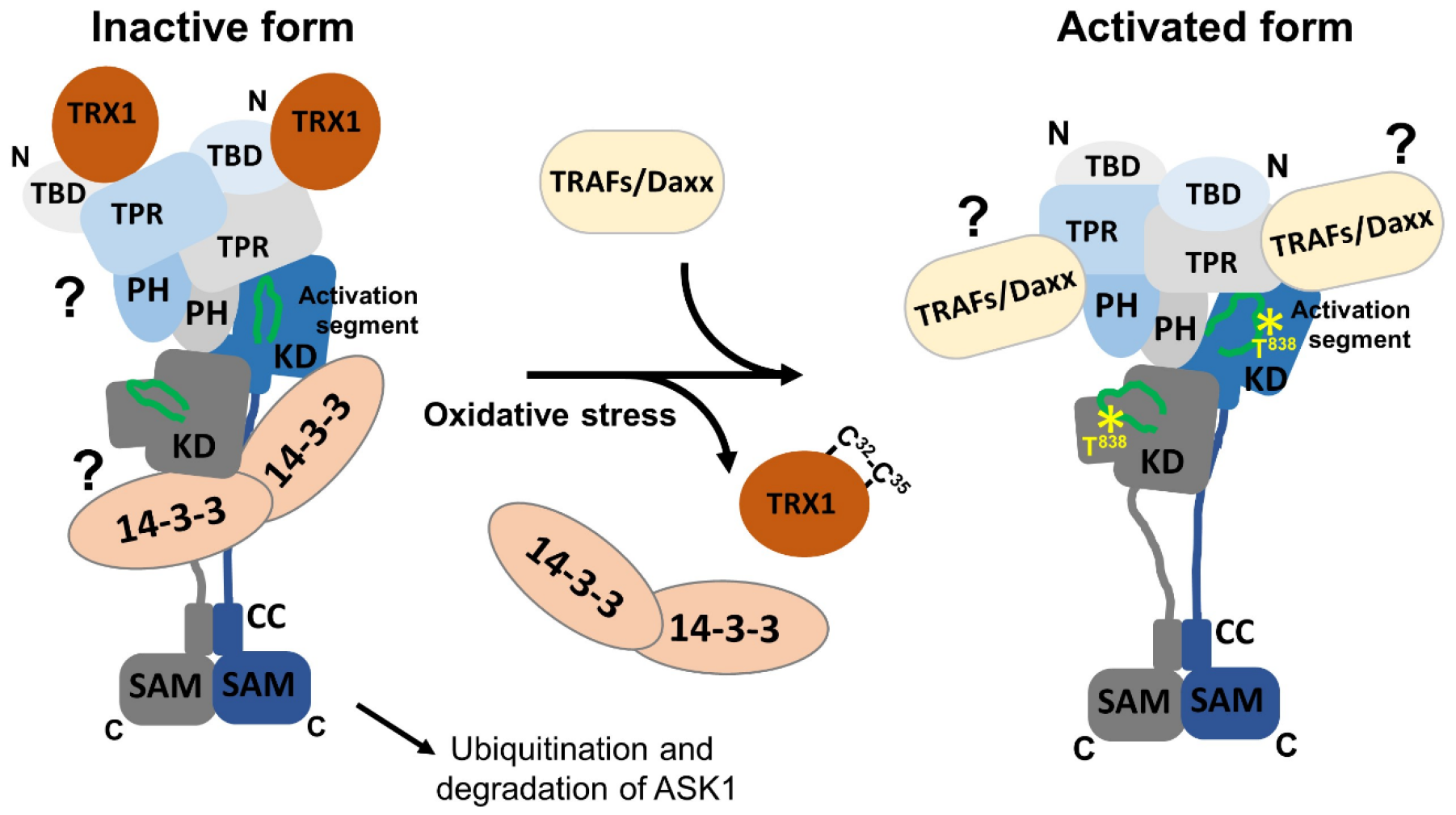
Proposed schematic model of ASK1 activation. In the resting state, ASK1 constitutively oligomerizes and is kept in an inactive state through interactions with TRX1 and 14-3-3 proteins. TRX1 appears to function as an allosteric effector whose binding affects the structure of TBD, likely affecting its interaction with TPR. Therefore, TRX1 affects the structure of CRR and allosterically modulates several regions of KD, even reducing access to the activation segment with the key phosphorylation site T838. Oxidative stress triggers TRX1 dissociation, followed by 14-3-3 dissociation and TRAF recruitment. These events subsequently lead to a conformational change in KD and increase access to the activation segment, enabling its phosphorylation at T838 and, as a result, stabilizing the active conformation. The role of 14-3-3 proteins in ASK1 inhibition and the mechanism whereby TRAFs and Daxx are involved in ASK1 activation remain unclear and should be explored in subsequent studies.

## Materials and Methods

### Recombinant protein expression and purification

Human TRX1 was expressed and purified as described previously^23^. To prevent TRX1 homodimerization caused by the formation of intermolecular disulfide bridges between non-active site C^73^ residues at high protein concentrations, we used the human TRX1 mutant C^73^S, which was reported to have unaltered structure and activity^51,52^.

DNA encoding the TBD domain of human ASK1 (residues 88–267) was ligated into pRSFDuet-1 (Merck KGaA, Darmstadt, Germany) using BamHI and NotI sites. Modified pRSFDuet-1 containing the sequence of the 6×His-tagged GB1 domain of protein G inserted into the first multiple cloning site was kindly provided by Evzen Boura (Institute of Organic Chemistry and Biochemistry AS CR, Prague, Czech Republic). ASK1 TBD was expressed at 25 °C for 18 h and purified from Escherichia coli BL21 (DE3) cells using Chelating Sepharose Fast Flow (GE Healthcare, Chicago, IL, USA) according to the standard protocol. The 6×His-tag was cleaved by incubation with TEV protease (250 U of TEV/mg of fusion protein) at 4 °C overnight during dialysis against a buffer containing 50 mM Tris-HCl (pH 8), 0.5 M NaCl, 4 mM EDTA, 80 mM imidazole, 4 mM 2-mercaptoethanol and 10% (w/v) glycerol. Chelating Sepharose Fast Flow (GE Healthcare, Chicago, IL, USA) was then used to capture the 6×His-GB1 and 6×His-TEV, and the flow-through sample containing the ASK1 TBD protein was purified by size-exclusion chromatography on a HiLoad 26/600 Superdex 75 pg column (GE Healthcare, Chicago, IL, USA) in a buffer containing 20 mM HEPES (pH 7.0), 200 mM NaCl, 5 mM EDTA, 20 mM glycine, 5 mM DTT and 10% (w/v) glycerol.

TBD-CRR, CRR, KD, TBD-CRR-KD and CRR-KD of human ASK1 (residues 88-658, 269-658, 658-973, 88-973 and 269-973, respectively) were ligated into the modified pRSFDuet-1 using BamHI and NotI sites or pST39 using XbaI and BamHI sites. Proteins were expressed at 18 °C (ASK1 CRR-KD and ASK1 TBD-CRR-KD) or 25 °C (ASK1 TBD-CRR and ASK1 CRR) for 18 h and purified from Escherichia coli BL21 (DE3) cells using Chelating Sepharose Fast Flow (GE Healthcare, Chicago, IL, USA) according to the standard protocol. Proteins were dialyzed overnight against a buffer containing 20 mM Tris-HCl (pH 7.5), 0.5 M NaCl, 1 mM EDTA, 2 mM 2-mercaptoethanol, and 10% (w/v) glycerol. ASK1 TBD-CRR and CRR constructs were incubated with TEV protease (250 U of TEV/mg of fusion protein) to remove 6×His-GB1 protein. ASK1 CRR-KD and TBD-CRR-KD contained uncleavable C-terminal 6×His-tag. The final purification step was size-exclusion chromatography on a HiLoad 26/600 Superdex 75/200 pg column (GE Healthcare, Chicago, IL, USA) in a buffer containing 20 mM Tris-HCl (pH 7.5), 150 mM NaCl, 5 mM DTT, and 10% (w/v) glycerol. ASK1 KD (residues 658-973) was expressed and purified as described previously^24^.

### Analytical ultracentrifugation

Sedimentation velocity (SV) experiments were performed using a ProteomLabTM XL-I analytical ultracentrifuge (Beckman Coulter, Brea, CA, USA). Samples were dialyzed against a buffer containing 20 mM Tris-HCl (pH 7.5), 150 mM NaCl and 2 mM 2-mercaptoethanol before the AUC measurements. SV experiments were conducted in charcoal-filled Epon centerpieces with a 12-mm optical path length at 20 °C and at rotor speeds ranging from 38,000 to 48,000 rpm (An-50 Ti rotor, Beckman Coulter). All sedimentation profiles were recorded with either absorption optics at 280 nm or interference optics. Buffer density and viscosity were estimated using the program SEDNTERP^53^. Diffusion-deconvoluted sedimentation coefficient distributions *c*(*s*) were calculated from raw data using the SEDFIT package^54^.

### Hydrogen/deuterium exchange coupled to mass spectrometry (HDX-MS)

ASK1 constructs (20 μM) and TRX1 were subjected to H/D exchange alone or in a mixture combining ASK1 TBD-CRR-KD with TRX1 (100 μM) and pre-incubated for 20 min at 4 °C. HDX reactions were performed by adding a 10× dilution of the protein mixture into a D_2_O-based buffer containing 20 mM Tris-HCl (pD 7.5), 200 mM NaCl, 2 mM 2-mercaptoethanol, and 10 % glycerol and by incubating at 4 °C. HDX was quenched after 2 s, 10 s, 1 min and 10 min of reaction by adding ice-chilled 1 M Glycine-HCl (pH 2.3), 6 M urea, 2 M thiourea, and 400 mM TCEP, in a 1:1 ratio, and all samples were frozen in liquid nitrogen. The 2 s and 10 min aliquots were prepared in triplicates. Thawed samples were loaded into the LC system including a custom-made pepsin/nepenthesin-2 protease column, and the generated peptides were online trapped and desalted on a SecurityGuard pre-column (ULTRA Cartridges UHPLC Fully Porous Polar C18, 2.1 mm, Phenomenex, Torrance, CA, USA) for 3 min under the flow of 0.4% formic acid (FA) in water, delivered at a flow rate of 200 μL.min^−1^ (1260 Infinity II Quaternary pump, Agilent Technologies, Waldbronn, Germany). Desalted peptides were then separated on a reversed-phase analytical column (LUNA Omega Polar C18 Column, 100 Å, 1.6 μm, 100 mm × 1.0 mm, Phenomenex, Torrance, CA, USA) at a flow rate of 40 μL.min^−1^ using a 10–40% linear gradient of solvent B (A: 2% acetonitrile/0.1% FA in water; B: 98% acetonitrile/0.1% FA in water) (1290 Infinity II LC system, Agilent Technologies, Waldbronn, Germany). The temperature within the customized LC system was kept at 0 °C to minimize the back exchange. All separated peptides were directly introduced into the ESI source of timsTOF Pro mass spectrometer with PASEF (Bruker Daltonics, Bremen, Germany). The data were analyzed using Data Analysis v. 5.3 (Bruker Daltonics, Bremen, Germany) and in-house DeutEx software. For each protein, peptides were identified by data-dependent LC–MS/MS using the same LC setup, performing a MASCOT (Matrix Science, London, UK) search against a custom-built database with sequences of ASK1, TRX1 and contaminants from the cRAP database.

### Cryo-EM - sample preparation and data collection

To prepare grids, thawed ASK1 TBD-CRR-KD was subjected to size exclusion chromatography on a Superdex 200 10/300 GL column (GE Healthcare, Chicago, IL, USA) in a buffer containing 20 mM Tris-HCl (pH 7.5), 150 mM NaCl and 2 mM 2-mercaptoethanol. The peak fraction of ASK1 TBD-CRR-KD was diluted in a 1:1 ratio with a buffer containing CHAPSO to a final concentration of 3.9 mM CHAPSO and a final concentration 0.9 mg/mL ASK1. Subsequently, 3.5 μL of protein solution was applied to a freshly glow-discharged (45 s total time, Gatan Solarus II 955 (Gatan, Inc., Pleasanton, CA, USA)) UltrAuFoil holey grids (R1.2/1.3, Quantifoil, Großlöbichau, Germany). Blotting was performed using a Vitrobot Mark IV, for 4 s, at 20°C and 100% humidity; all grids were plunge-frozen in liquid ethane and stored in liquid nitrogen until use. The grids were screened under a JEOL JEM 2100-plus electron microscope (Akishima, Tokyo, Japan) at 200 keV equipped with a TVIPS TemCam– XF416 4K CMOS camera (TVIPS GmbH, Gauting, Germany) and under a Talos Arctica electron microscope (FEI, Thermo Fisher Scientific, Hillsboro, Oregon, USA) at 200 keV equipped with a GATAN K2 Summit detector (Gatan, Inc., Pleasanton, CA, USA). All data were collected under a Titan Krios electron microscope (FEI, Thermo Fisher Scientific, Hillsboro, Oregon, USA), at 300 keV, equipped with a Gatan K3 BioQuantum detector (Gatan, Inc., Pleasanton, CA, USA). Movies were recorded at 105,000× magnification and 0.834 Å per pixel calibrated resolution. The defocus values ranged from −0.7 to −2.8 μm, with a total exposure of 40 e^−^/Å^2^. From 11395 movies, 8691 were collected under a 40° tilt. Each movie consisted of 40 frames.

### Cryo-EM - image processing

All images were processed in CryoSPARC 4.1.2^55^. Once the movies were imported and gain-corrected, they were subjected to patch motion correction and patch CTF estimation. Micrographs with an estimated CTF resolution > 5 Å, full-frame motion distance > 20 Å and a relative ice thickness > 1.2 were discarded; then, micrographs were visually curated, and those with excessive aggregation, ice artifacts or artifacts in power spectra were also excluded. The initial particle set was picked from 200 randomly selected micrographs, using a blob picker tool with 60-180 Å particle size. After visual inspection, particles were extracted with a box size of 320×320 pixels, and after 2D classification, good classes were used to generate templates. Particle picking was then repeated using a template based picker. Particle picks were extracted with a box size of 340×340 pixels and subjected to 2D classification. Particles within good 2D classes were used for ab-initio 3D reconstruction, heterogeneous refinement, and homogeneous and non-uniform refinements with separated classes, including the calculation of gold standard Fourier shell correlation (GSFSC). GSFSC was used to determine the final map resolution with 0.143 FSC threshold. The resulting map was sharpened using phenix.autosharpen^56^. Details about data processing workflow are provided in Supplementary Table S1 and Fig. S1.

### Cryo-EM - model building, refinement and analysis

After visual inspection of the final map, we used the “jiggle-fit” tool in Coot 0.9.8^57^ to fit known crystal structures of different ASK1 domains, namely KD (PDB ID: 2CLQ^13^) and ASK1 CRR (PDB ID: 5ULM ^14^). Then, we fitted the Alphafold prediction (AF-Q99683-F1) to place a model for TBD. After finishing connections and adjustments, the model was excluded in areas of insufficient or uninterpretable density. Atomic refinement was performed using phenix.real_space_refine from the Phenix 1.19.1 software package^56^. The model was validated using MolProbity^58^. Model statistics are presented in Supplementary Table S1. The final model has been deposited in PDB/EMDB under accession code: 8QGY/EMD-18396.

## Data availability

The authors declare that all data supporting the findings of this study are available within the article and its supplementary information file. Cryo-EM data have been deposited in the RCSB PDB/EMDB with the accession code: 8QGY/EMD-18396.

## Acknowledgments

This study was funded by the Czech Science Foundation (grant No. 19-00121S), the Grant Agency of the Charles University (K.H., grant number 1160120) and the Czech Academy of Sciences (RVO: 67985823 of the Institute of Physiology). We acknowledge CMS-Biocev (“Biophysical techniques, Crystallization, Diffraction, Structural mass spectrometry”) of CIISB and Cryo-electron microscopy and tomography core facility CEITEC MU of CIISB, Instruct-CZ Centres, supported by MEYS CR (LM2018127) and CZ.02.1.01/0.0/0.0/18_046/0015974. We thank P. Pompach and P. Vankova for their help with MS measurements, J. Miksatko, J. Novacek and M. Pinkas for their assistance with cryo-EM data collection and Carlos V. Melo for editing the article.

