## Supporting information for "The cryo-EM structure of ASK1 reveals an asymmetric architecture allosterically modulated by TRX1"

### Contents

**1. Supporting Tables**
1.1 Table S1: Cryo-EM data collection, refinement and validation statistics 2

**2. Supporting Figures**
2.1 Figure S1: ASK1 TBD-CRR-KD data processing 3
2.2 Figure S2: ASK1 TBD-CRR-KD map quality assessment 4
2.3 Figure S3: Agreement between the experimental density map and the refined model of C-terminally truncated ASK1 5
2.4 Figure S4: Secondary structure of ASK1 TBD-CRR-KD 6
2.5 Figure S5: Comparison of CRR regions with the crystal structure of the isolated CRR 7
2.6 Figure S6: Comparison of KDs with the crystal structure of the isolated KD 8
2.7 Figure S7: SV AUC analysis showing the effect of TRX1 binding on the dimerization of ASK1 TBD-CRR-KD 9
2.8 Figure S8: Sequence coverage of ASK1 TBD-CRR-KD (residues 88−973) assessed by HDX 10
2.9 Figure S9: TRX1 binding induces structural changes in all domains of ASK1 TBD-CRR-KD 11
2.10 Figure S10: Effect of TRX1 binding on TBD deuteration in the ASK1 TBD-CRR-KD dimer 12
2.11 Figure S11: Effect of TRX1 binding on CRR deuteration in the ASK1 TBD-CRR-KD dimer 13
2.12 Figure S12: Effect of TRX1 binding on KD deuteration in the ASK1 TBD-CRR-KD dimer 14
2.13 Figure S13: Sequence coverage of TRX1 assessed by HDX 15
2.14 Figure S14: Effect of complex formation on TRX1 deuteration 16

**3. References** 19

**Table S1. Cryo-EM data collection, refinement and validation statistics**

| | |
|---|---|
| **Data collection and image processing** | |
| Instrument | FEI Titan Krios |
| Camera | K3 Gatan |
| Magnification | 105,000 |
| Voltage (kV) | 300 |
| Electron exposure ($e^-/Å^2$) | 40 |
| Defocus range (μm) | −0.7 to −2.8 |
| Pixel size (Å) | 0.834 |
| Total number of micrographs 0°tilt (no.) | 2704 |
| Total number of micrographs 40°tilt (no.) | 8691 |
| Selected micrographs (mixed, no.) | 5781 |
| Initial particle images | 1126189 |
| Final particle images (mixed, no.) | 645915 |
| Starting model | Ab initio |
| Symmetry | C1 |
| Map resolution, masked (Å)/FSC threshold | 3.71/0.143 |
| Sharpening | phenix.autosharpen |
| EMDB code | EMD-18396 |
| **Model building and refinement** | |
| Initial models used, PDB or AlphaFold Protein Structure database codes: | 2CLQ (KD)<br>5ULM (CRR)<br>AF-Q99683-F1 (TBD) |
| Masked FSC (map-model)/FSC threshold | 3.66/0.143 |
| Non-hydrogen atoms | 12744 |
| Amino acid residues | 1589 |
| Protein molecules | 2 |
| Real-space correlation - CCvolume | 0.64 |
| Real-space correlation - CCmasked | 0.64 |
| B factors (mean, $Å^2$) | 97.92 |
| Root-mean-square deviation of bond length (Å)/angles (°) | 0.003/0.616 |
| PDB code | 8QGY |
| **Validation** | |
| MolProbity score | 2.09 |
| Clashscore | 4.56 |
| Rotamer outliers (%) | 3.9 |
| Ramachandran plot - Favored (%)/Allowed (%)/Disallowed (%) | 93.77/6.17/0.06 |

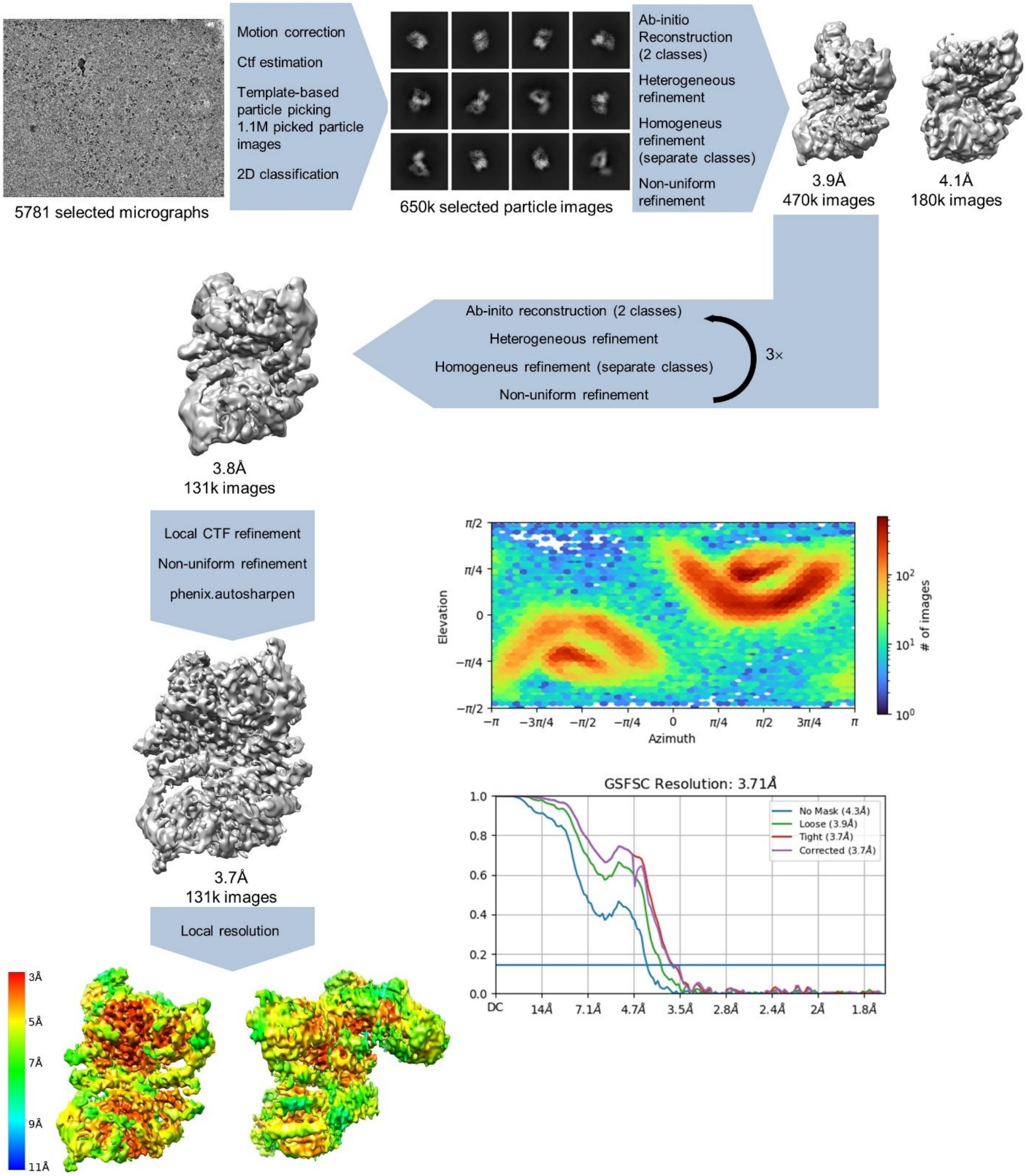

**Figure S1. ASK1 TBD-CRR-KD data processing.** Processing steps are indicated in the scheme, showing the number of particle images, resolution, CryoSPARC 4.1.2 output for GSFSC analysis and view distribution plots. The maps were calculated using either the threshold 0.05-0.1 or the threshold 5 for sharpened maps.

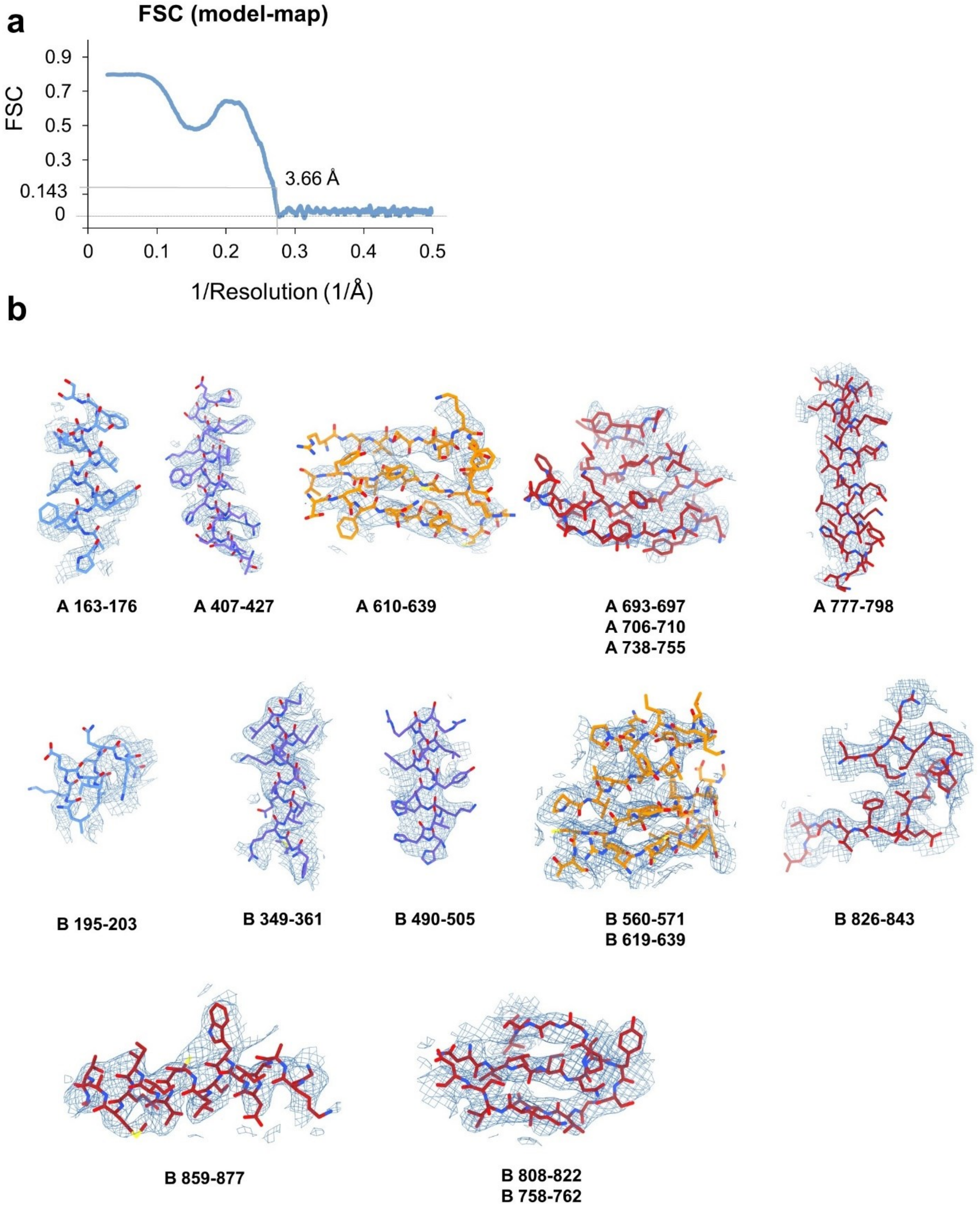

**Figure S2. ASK1 TBD-CRR-KD map quality assessment.** (a) FSC (model-map) reported by the Phenix validation tool (1), indicating the threshold 0.143. (b) Examples of the final density map fitted to the model. The maps were generated in ChimeraX (2) using a threshold level ranging from 7 to 13.

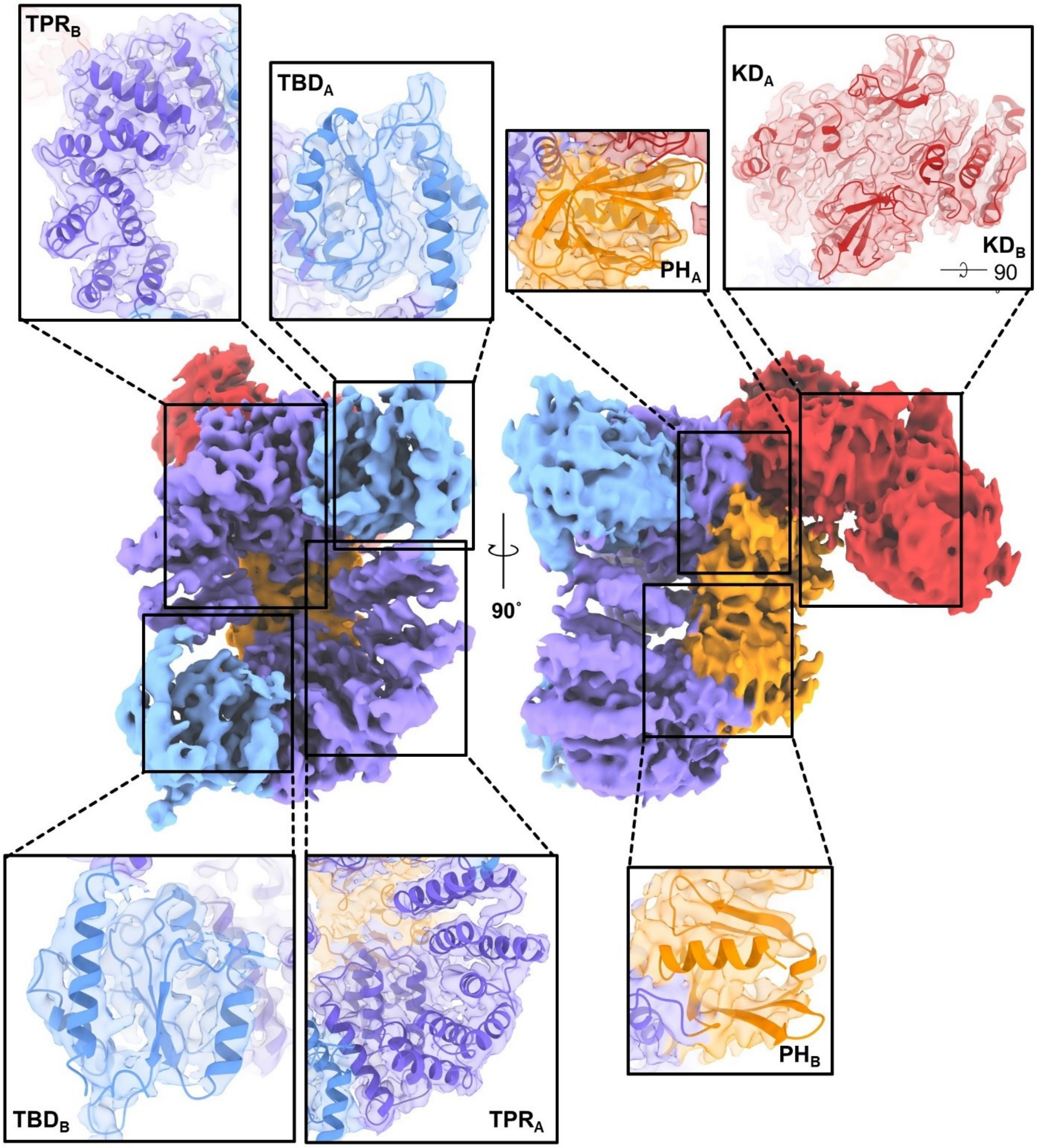

**Figure S3. Agreement between the experimental density map and the refined model of C-terminally truncated ASK1.** The maps were generated using a threshold level of 5-7 and 80% transparency.

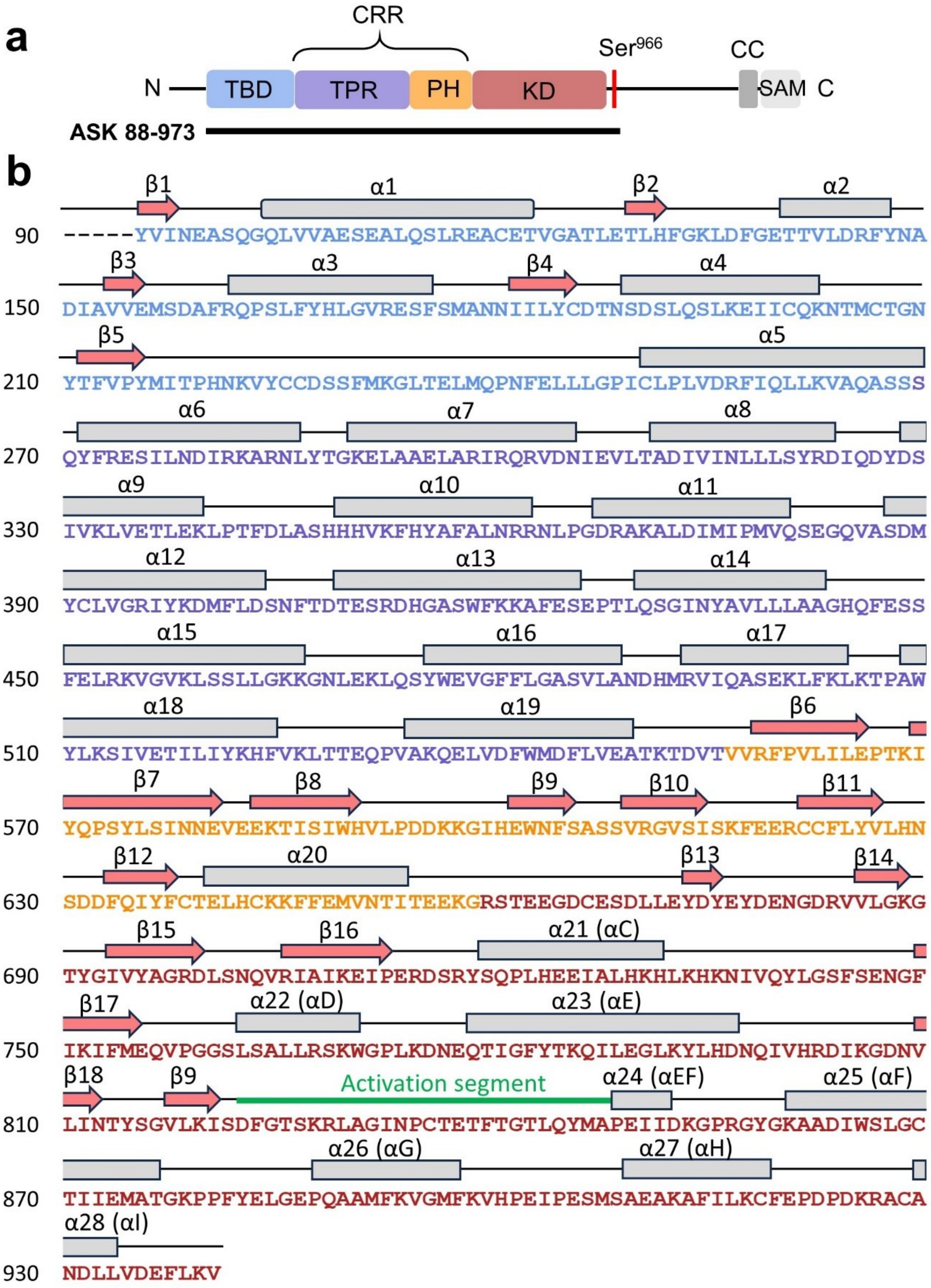

**Figure S4. Secondary structure of ASK1 TBD-CRR-KD.** (a) Domain structure of ASK1. The black line indicates the construct used for cryo-EM structural analysis. (b) Secondary structure of the cryo-EM model of ASK1 TBD-CRR-KD.

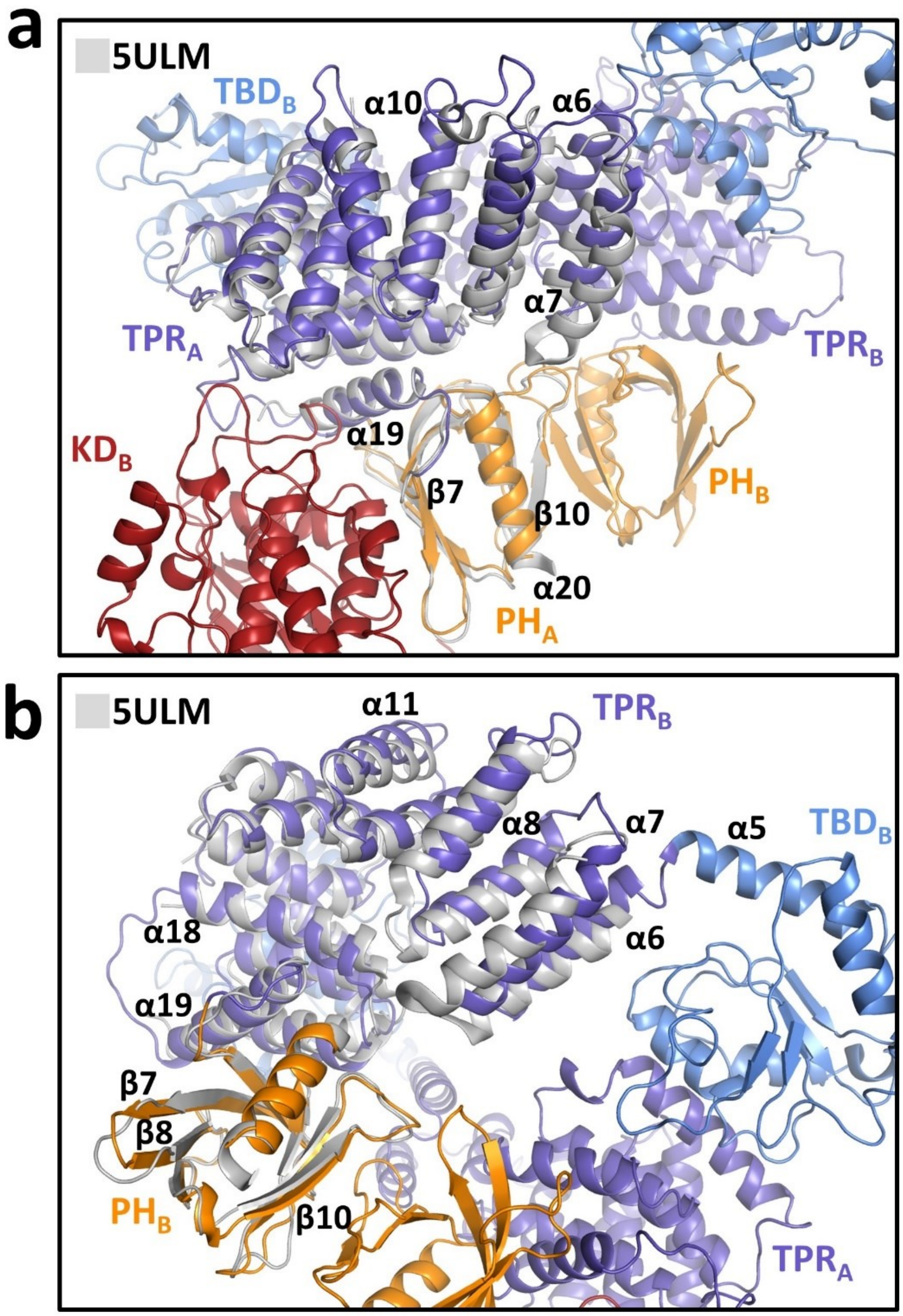

**Figure S5. Comparison of CRR regions with the crystal structure of the isolated CRR.** Superposition of the X-ray structure of CRR (PDB ID: 5ULM (3), shown in gray) and $CRR_A$ (a) and $CRR_B$ (b) from the cryo-EM model of ASK1 TBD-CRR-KD. TPR, tetratricopeptide repeats; PH, pleckstrin-homology domain; KD, kinase domain.

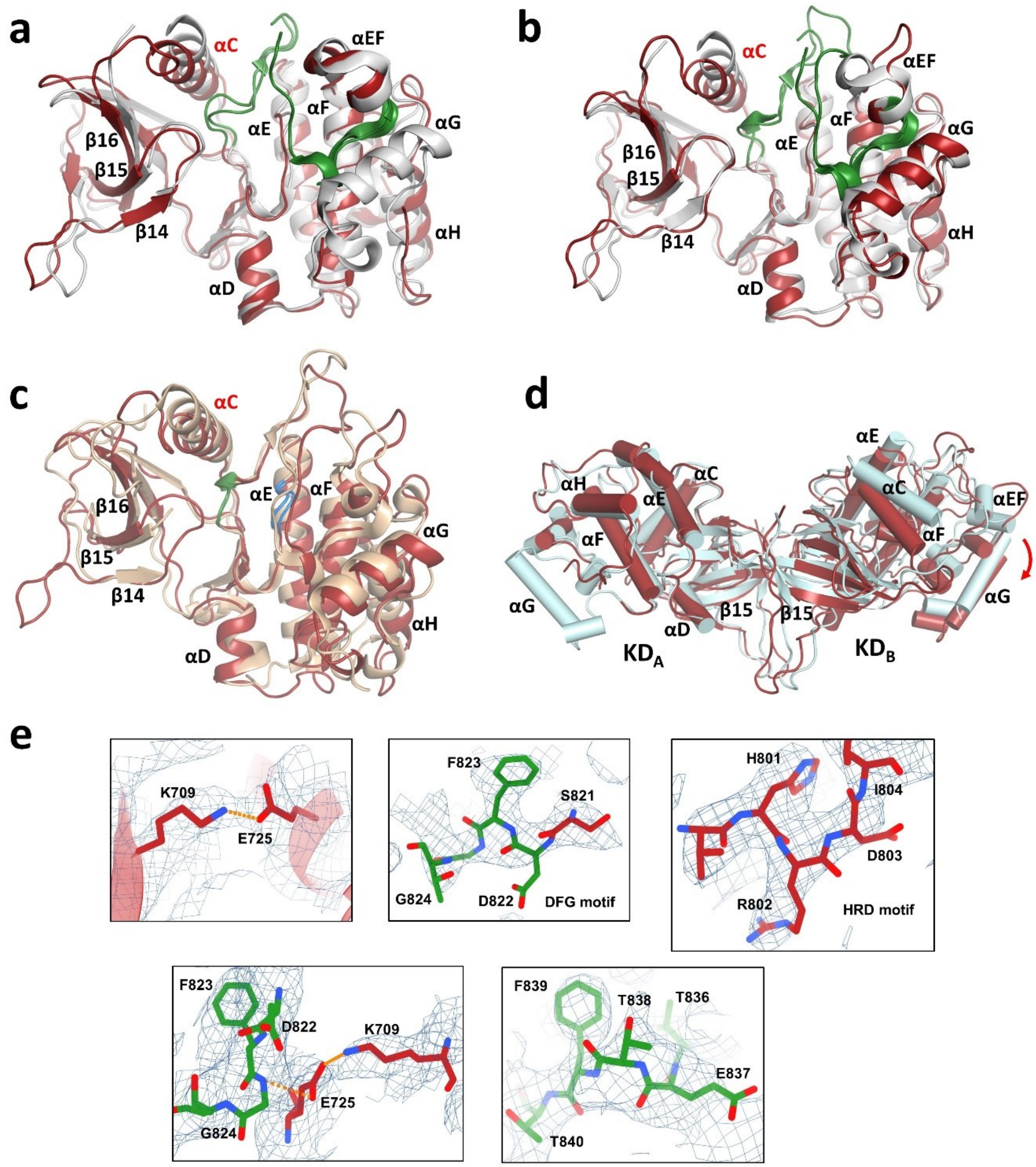

**Figure S6. Comparison of KDs with the crystal structure of the isolated KD.** (a) Superposition of the X-ray structure of KD with bound inhibitor (PDB ID: 2CLQ (4), shown in grey) and $KD_A$ from the cryo-EM model of ASK1 TBD-CRR-KD (shown in red). The activation segment is shown in green. (b) Superposition of 2CLQ (grey) and $KD_B$ (red). (c) Superposition of the X-ray structure of BRAF (PDB ID: 4MNE (5), shown in sand) and $KD_B$ (red). The DFG motif is shown in green, the HRD motif is shown in blue. (d) Cryo-EM structure of KD dimer (red) superimposed with the x-ray structure of KD dimer (cyan, PDB ID: 2CLQ) via $KD_A$. Helices are represented as cylinders. The red arrow indicates the shift of the C-lobe of the $KD_B$. (e) Representing density of selected fragments of $KD_B$. Residues and/or visualized motives are indicated in panels. The maps were calculated using a threshold level of 8-15.

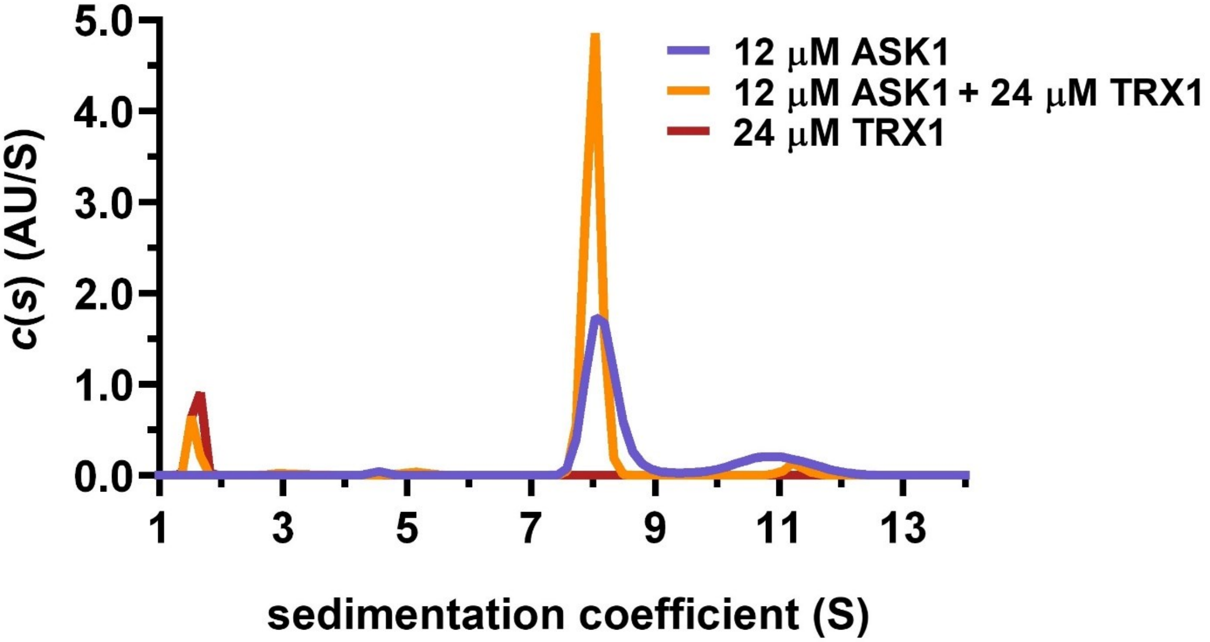

**Figure S7. SV AUC analysis showing the effect of TRX1 binding on the dimerization of ASK1 TBD-CRR-KD.** Distribution of sedimentation coefficients, $c(s)$, of 12 μM ASK1 TBD-CRR-KD alone, 12 μM ASK1 TBD-CRR-KD with 24 μM TRX1 and 24 μM TRX1 alone are shown in violet, orange and brown, respectively.

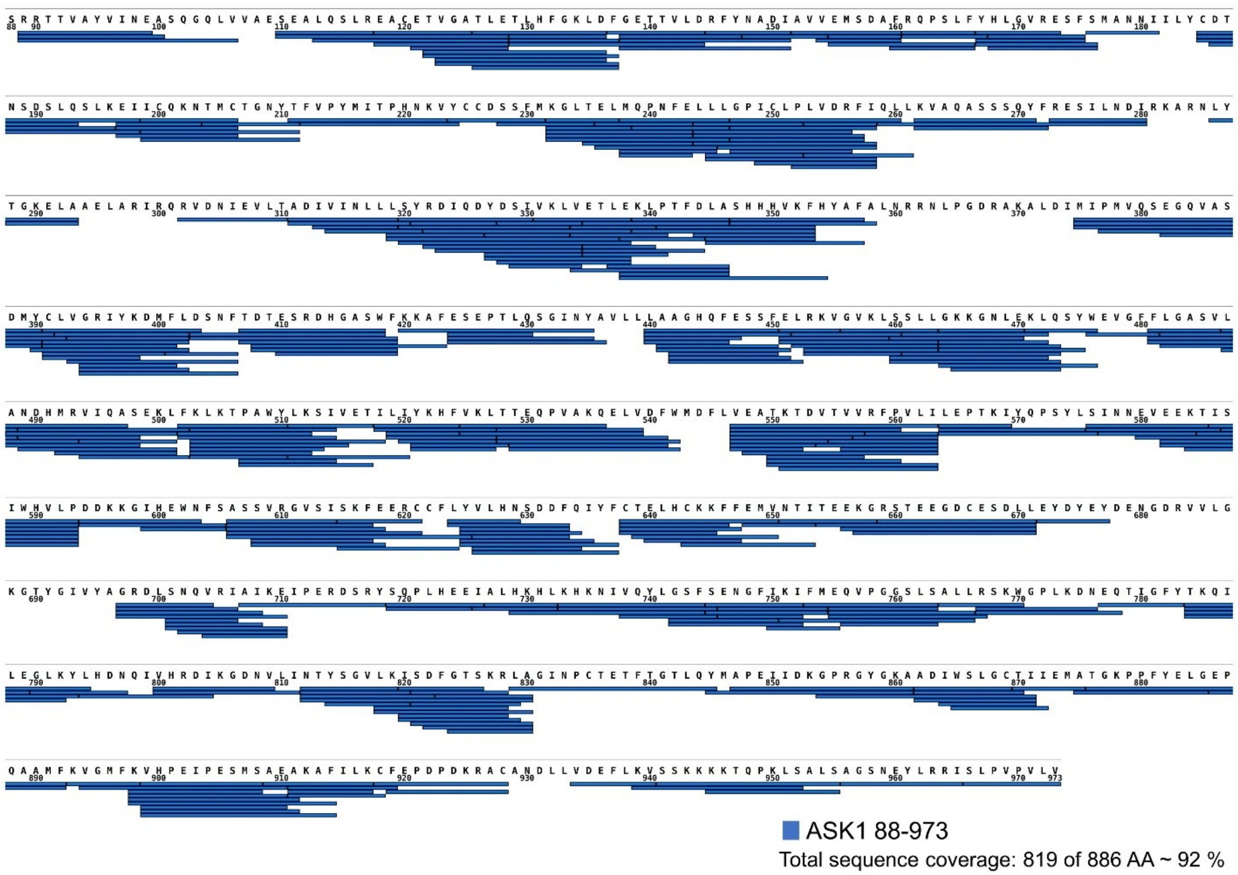

**Figure S8. Sequence coverage of ASK1 TBD-CRR-KD (residues 88−973) assessed by HDX.** The sequence coverage of the construct reached 92%, with 819 of 886 amino acid residues. The map was created using the DrawMap script of MSTools (http://peterslab.org/MSTools/).

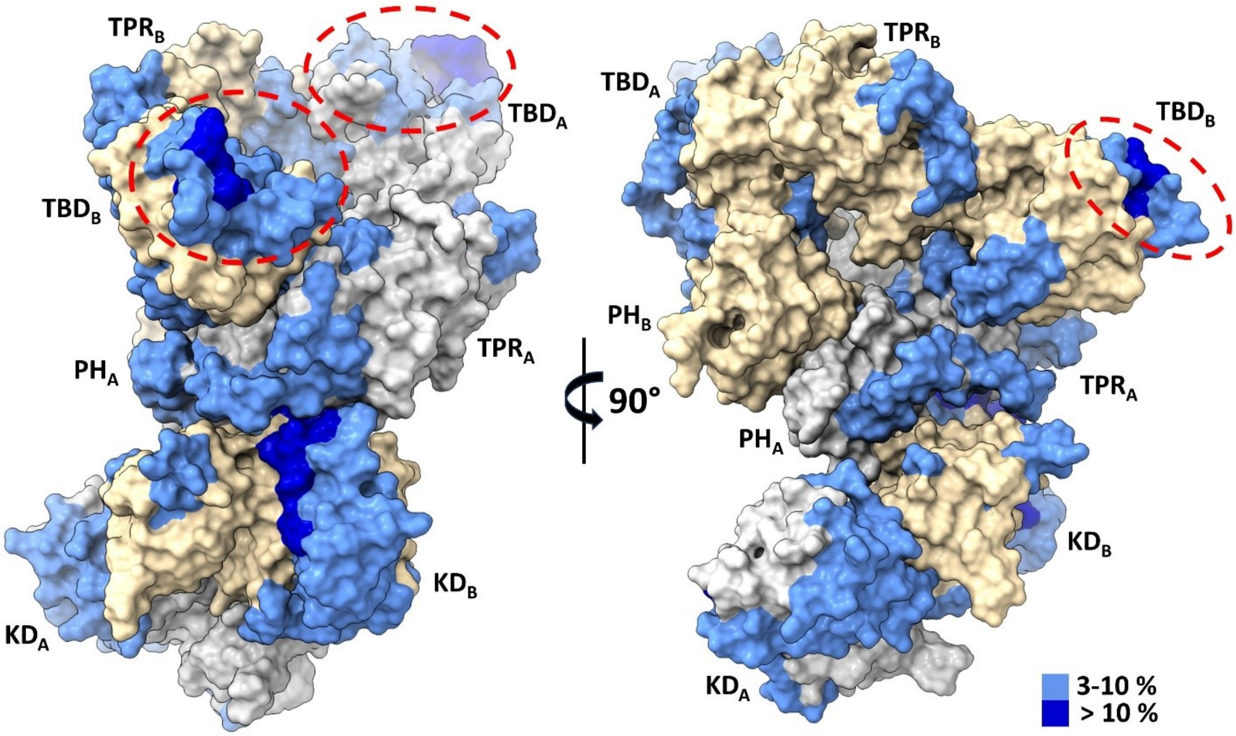

**Figure S9. TRX1 binding induces structural changes in all domains of ASK1 TBD-CRR-KD.** (a) Surface representation of the structure of the ASK1 TBD-CRR-KD dimer (chain A in grey, chain B in light yellow) colored according to changes in deuteration in the presence of TRX1 after 600 s. Red ellipses indicate TRX1-binding sites on TBDs.

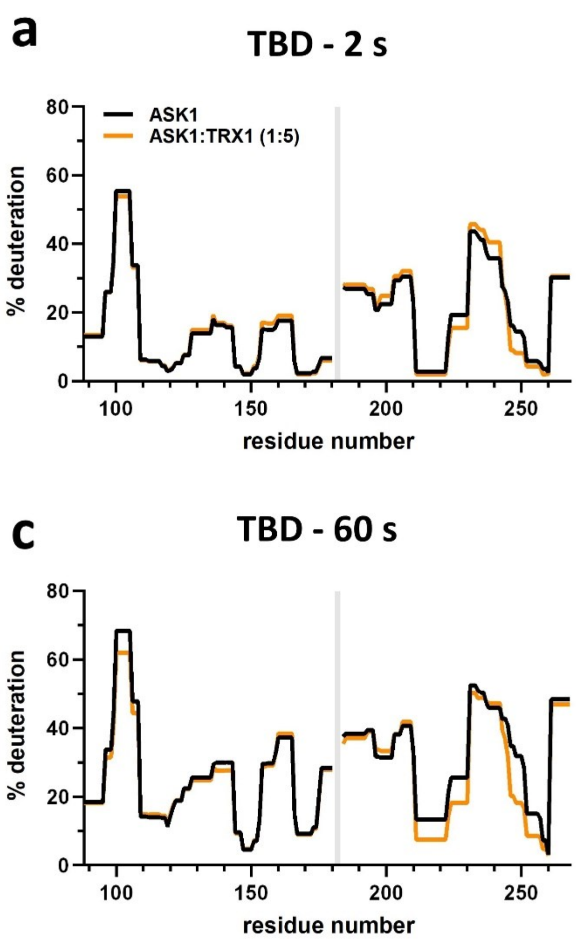

**Figure S10. Effect of TRX1 binding on TBD deuteration in the ASK1 TBD-CRR-KD dimer.** Protection plots showing differences in TBD deuteration within ASK1 TBD-CRR-KD with (orange) or without (black) TRX1 at four different deuteration times: 2 s, 10 s, 1 min and 10 min. Grey zones indicate areas without coverage.

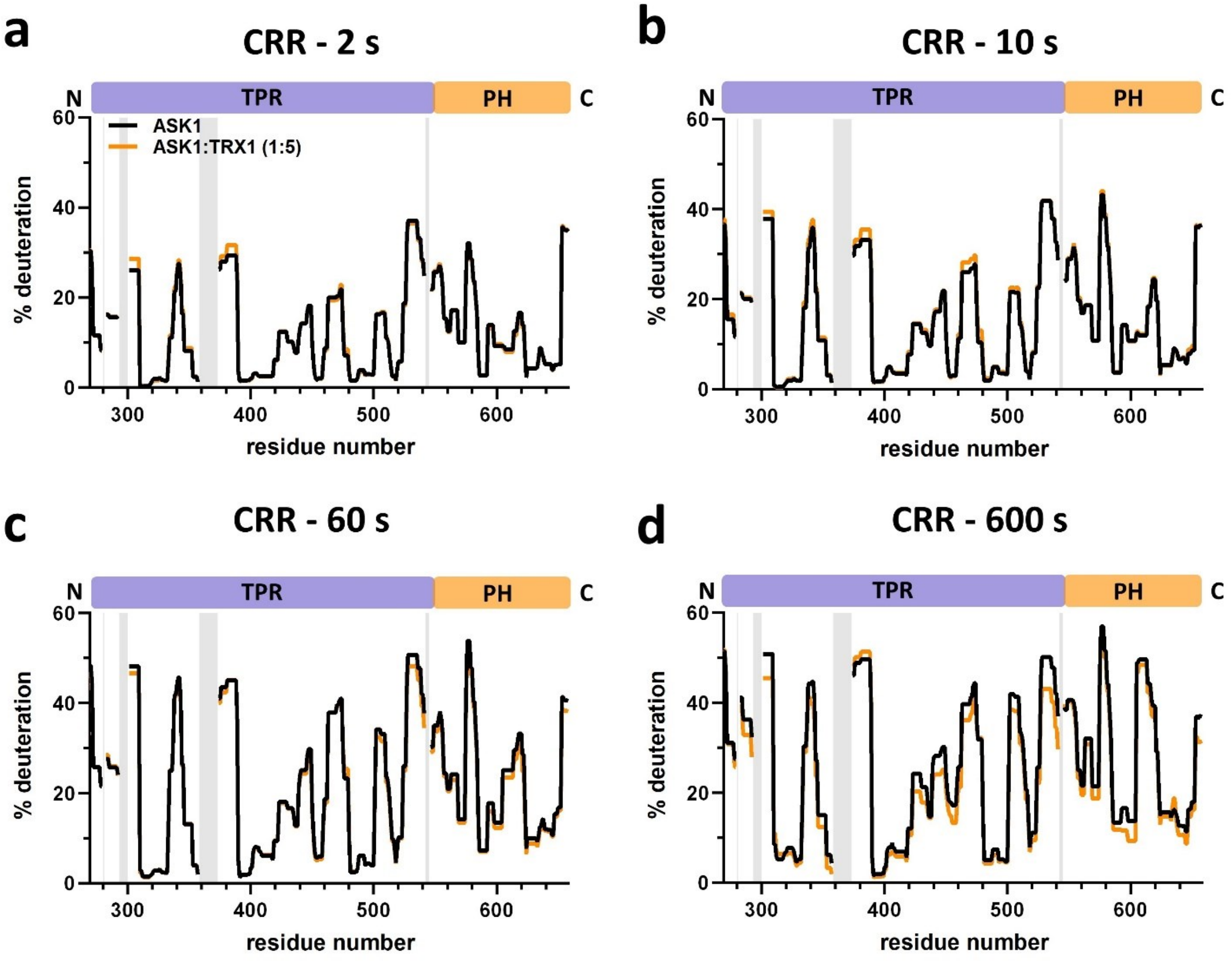

**Figure S11. Effect of TRX1 binding on CRR deuteration in the ASK1 TBD-CRR-KD dimer.** Protection plots showing differences in CRR deuteration within ASK1 TBD-CRR-KD with (orange) or without (black) TRX1 at four different deuteration times: 2 s, 10 s, 1 min and 10 min. Grey zones indicate areas without coverage.

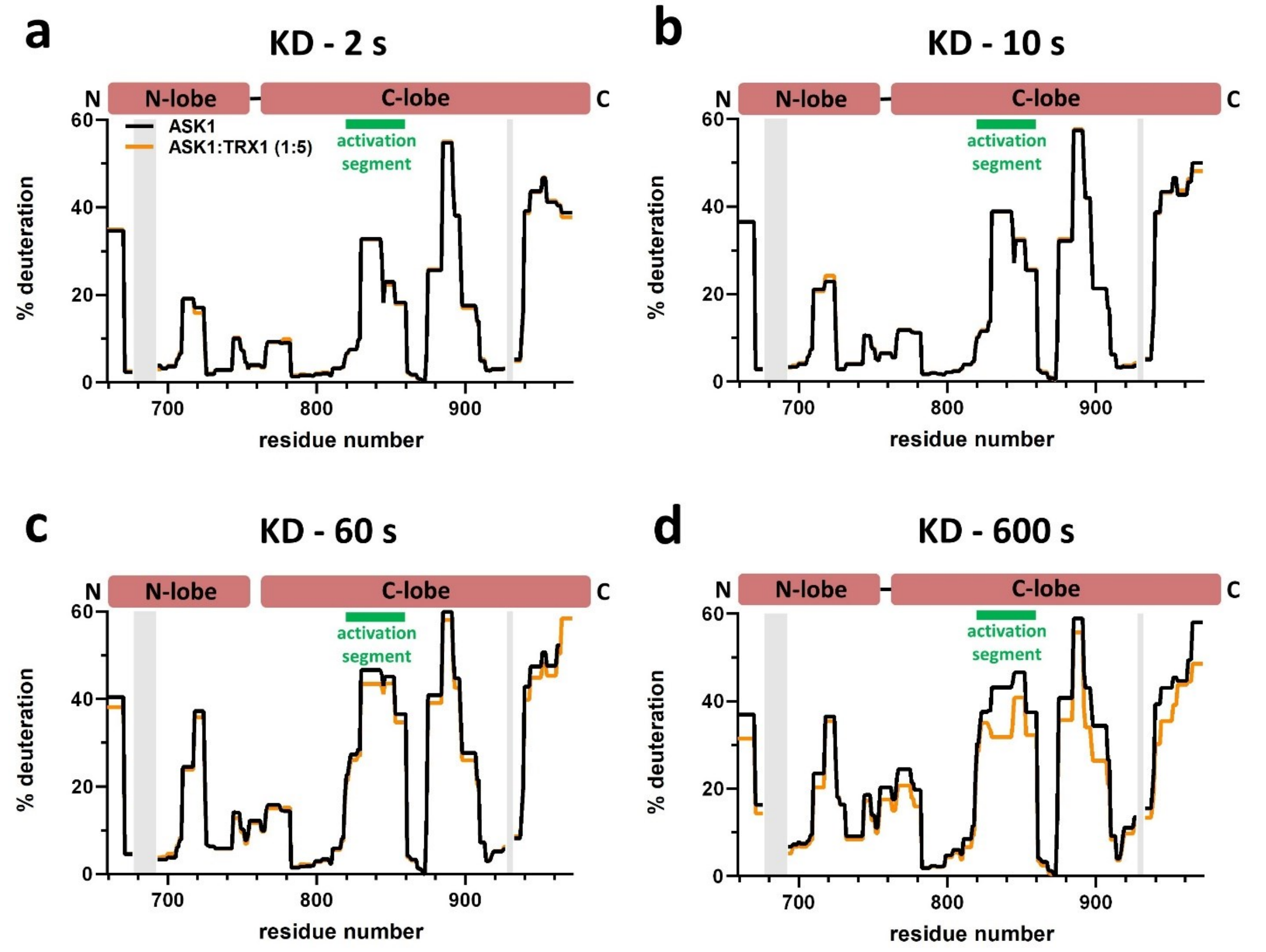

**Figure S12. Effect of TRX1 binding on KD deuteration in the ASK1 TBD-CRR-KD dimer.** Protection plots showing differences in KD deuteration within ASK1 TBD-CRR-KD with (orange) or without (black) TRX1 at four different deuteration times: 2 s, 10 s, 1 min and 10 min. Grey zones indicate areas without coverage.

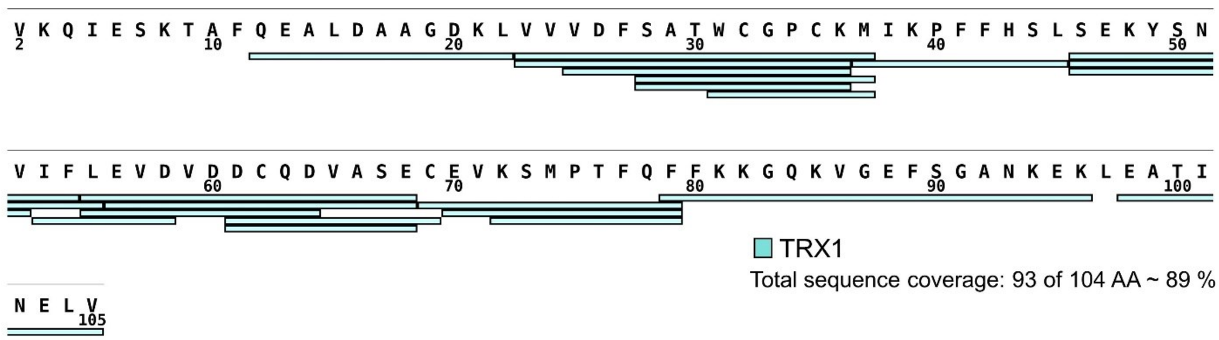

**Figure S13. Sequence coverage of TRX1 assessed by HDX.** The sequence coverage of the TRX1 construct reached 89%, with 93 of 104 amino acid residues. The map was created using the DrawMap script of MSTools (http://peterslab.org/MSTools/).

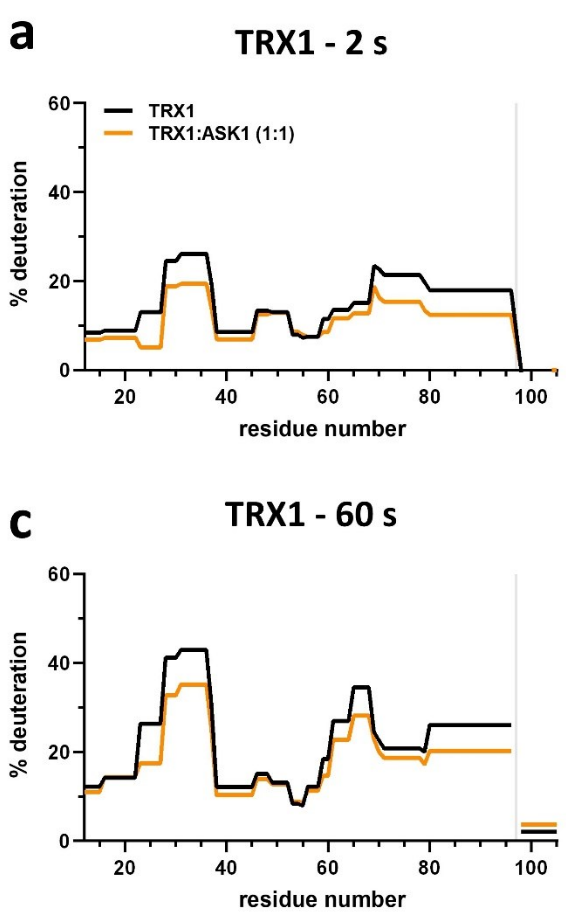

**Figure S14. Effect of complex formation on TRX1 deuteration.** Protection plots showing differences in KD deuteration within ASK1 TBD-CRR-KD with (orange) or without (black) TRX1 at four different deuteration times: 2 s, 10 s, 1 min and 10 min. Grey zones indicate areas without coverage.
